# Comprehensive deletion landscape of CRISPR-Cas9 identifies minimal RNA-guided DNA-binding modules

**DOI:** 10.1101/2020.10.19.344077

**Authors:** Arik Shams, Sean A. Higgins, Christof Fellmann, Thomas G. Laughlin, Benjamin L. Oakes, Rachel Lew, Maria Lukarska, Madeline Arnold, Brett T. Staahl, Jennifer A. Doudna, David F. Savage

## Abstract

Proteins evolve through the modular rearrangement of elements known as domains. It is hypothesized that extant, multidomain proteins are the result of domain accretion, but there has been limited experimental validation of this idea. Here, we introduce a technique for genetic <u>m</u>inimization by <u>i</u>terative <u>s</u>ize-<u>e</u>xclusion and <u>r</u>ecombination (MISER) that comprehensively assays all possible deletions of a protein. Using MISER, we generated a deletion landscape for the CRISPR protein Cas9. We found that Cas9 can tolerate large single deletions to the REC2, REC3, HNH, and RuvC domains, while still functioning *in vitro* and *in vivo*, and that these deletions can be stacked together to engineer minimal, DNA-binding effector proteins. In total, our results demonstrate that extant proteins retain significant modularity from the accretion process and, as genetic size is a major limitation for viral delivery systems, establish a general technique to improve genome editing and gene therapy-based therapeutics.

## Introduction

Domains are the fundamental unit of protein structure ^1–3^. Domains are also the unit of evolution in proteins, accumulating incremental mutations that change their function and stability, as well as being recombined within genomes to create new proteins via insertions, fusions, or deletions ^4–7^. Extant multidomain proteins are thus thought to have evolved via the continuous accretion of domains to gain new function ^4 8,9^. Additionally, eukaryotic proteome diversity is vastly increased by alternative splicing, which tends to insert or delete protein domains ^10^. The phenomenon of domain modularity in proteins has been exploited synthetically to rearrange and expand the architecture of a protein, enabling new functionality ^11,12^. For example, the programmable DNA nuclease Cas9 can be converted into a ligand-dependent allosteric switch using advanced molecular cloning, similar to other domain insertions dictated by allostery ^13^. Although there are several methods for comprehensively altering protein topology ^14–16^, no method has been demonstrated for domain deletion.

Rationally constructed protein deletions have long been essential to elucidating functional and biochemical properties but are generally limited to a handful of truncations. Moreover, protein engineering can make use of deletions to alter enzyme substrate specificity ^17^, enable screens for improved activity and thermostability ^18^, or minimize protein size ^19^. Early approaches to protein deletion libraries resulted in the deletion of single amino acids using an engineered transposon ^20,21^. Other methods utilize direct PCR ^22^, random nuclease digestion ^23^, or random *in vitro* transposition followed by a complicated cloning scheme ^24^ to achieve deletion libraries containing a variety of lengths. These techniques are low in throughput and/or require complex molecular techniques which poorly capture library diversity - in contrast to protein insertions where library size grows linearly with target length, deletion libraries grow as the square.

A simple and efficient method for building protein deletions coupled with a selection strategy would provide the ability to comprehensively query and delineate the function of domains or motifs in complex and multi-domain proteins. Such a technique could be used to identify crucial functions within multidomain proteins or splicing variants in a manner akin to how deep mutational scanning can be used to identify the effects of single nucleotide polymorphisms on functionality ^25^. Moreover, with sufficient modularity, the evolutionary path of domain accretion could be explored through iterative combining, or ‘stacking,’ of domain deletions to isolate a minimal, core protein for a defined function ^9^.

One attractive target for such a strategy is *Streptococcus pyogenes* Cas9 (SpCas9), the prototypal RNA-guided DNA endonuclease used for genome editing ^26^. SpCas9 is an excellent model protein for a comprehensive deletion study because of its multi-domain architecture and availability of high-throughput assays for either DNA cutting or binding ^27^. Functionally, SpCas9 targets and cleaves DNA in a multi-step process. First, an apo Cas9 molecule forms a complex with a guide RNA (gRNA), containing a 19-22 bp variable “spacer” sequence that is complementary to a DNA target locus. The primed ribonucleoprotein (RNP) complex then surveills genomic DNA for a protospacer-adjacent motif (PAM) - 5’-NGG-3’ in the case of SpCas9, where N is any nucleobase - that initiates a transient interaction with the protein to search for an adjacent ~20-bp target sequence. If a target is present, the double-stranded DNA (dsDNA) helix is unwound, allowing the gRNA to anneal to the DNA and form a stable RNA-DNA hybrid structure called an R-loop (see illustration in Fig S8). Formation of a complete 20-bp R-loop triggers a conformational change in Cas9 to form the catalytically-active complex ^28–30^.

SpCas9 has a bi-lobed architecture consisting of the <u>REC</u>ognition lobe, responsible for recognizing and binding DNA sequences, and the <u>NUC</u>lease lobe, which possesses HNH and RuvC domains that cut the target and non-target strands of DNA, respectively. Cas9 is postulated to have evolved via domain accretion from a progenitor RuvC domain ^31 9^. As a consequence, Cas9 orthologs possess manifold architectures. For example, the SpCas9 REC lobe possesses three domains (REC1, REC2 and REC3) while the *Staphylococcus aureus* Cas9 (SaCas9) has a contiguous REC domain without REC2 ^32,33^. The function of REC2 is ambiguous but is thought to act as a conformational switch to trigger DNA cleavage ^34,35^, raising the question of how SaCas9 accomplishes the effect ^36^. Thus, the multi-domain, multi-functional nature of Cas9s make them an excellent model system for exploring domain deletions. Relatedly, Cas9’s large size also complicates its delivery using viral vectors. Knowledge of functional deletions may thus facilitate the delivery of genome editing therapeutics.

Here, we introduce genetic <u>m</u>inimization by <u>i</u>terative <u>s</u>ize-<u>e</u>xclusion and <u>r</u>ecombination (MISER), a technique for systematically exploring deletions within a protein. Application of MISER to SpCas9 identified regions of the protein which can be deleted with minimal consequence. Furthermore, we stacked individual deletions to engineer novel CRISPR Effector (CE) proteins that are less than 1000 amino acids in length. CRISPRi and biochemical assays demonstrated that these variants remain competent for target DNA-binding but are less functional than single deletion variants. Finally, to understand the structural consequence of deletion, we used single-particle cryo-electron microscopy to solve a 6.2 Å structure of the smallest, 874 amino acid CE. This structure surprisingly revealed an overall conformation that preserves essential functions of SpCas9—emphasizing the concept of domains as independent modules—even though the quaternary structure is severely modified.

## Results

### MISER reveals the comprehensive deletion landscape of SpCas9

The general concept of MISER is to create a pool of all possible contiguous deletions of a protein and analyze them in a high-throughput fitness assay. The process can then be iterated to stack deletions together. We created such a library by: i) programmably introducing two distinct restriction enzyme sites, each once, across a gene on an episomal plasmid, ii) excising the intervening sequence using the restriction enzymes and iii) re-ligating the resulting fragments (Fig. 1A). In the instantiation here, two separate restriction enzymes (NheI and SpeI) with compatible sticky ends are used. Cleavage, removal of intervening sequence, and ligation thus results in a two-codon scar (encoding either Ala-Ser or Thr-Ser) site not recognized by either enzyme, thereby increasing efficiency of cloning and enabling iteration of the entire process (Fig. S1).

**Figure 1:**
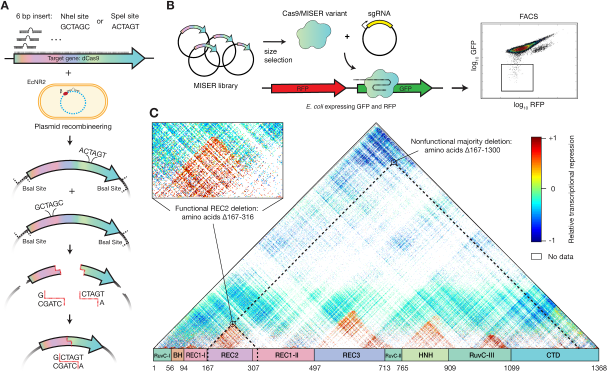
Minimization by Iterative Size Exclusion and Recombination (MISER). **A)** MISER library construction. A 6-bp SpeI or NheI recognition site is inserted separately into a dCas9-encoding plasmid flanked by BsaI sites using plasmid recombineering. The resultant libraries are digested with BsaI and either SpeI or NheI, and the two fragment pools are combined and ligated together to generate a library of dCas9 ORFs possessing all possible deletions. **B)** The MISER library is cloned into a vector and co-transformed in *E. coli* expressing RFP and GFP with an sgRNA targeting GFP. The library products are expressed, functional variants bind to the target, and repress the fluorophore. Repression activity *in vivo* is measured by flow cytometry. **C)** Enrichment map of the MISER deletion landscape of *S. pyogenes* dCas9. A single pixel within the map represents an individual variant that contains a deletion beginning where it intersects with the horizontal axis moving to the left (N) and ends where it intersects with the axis moving to the right (C). Larger deletions are at the top, with some deletions almost spanning the whole protein. The heatmap shows relative repression activity of variants from two FACS sorts of a single replicate.

The MISER library was made for nuclease-dead Cas9 (dCas9) as follows. First single NheI or SpeI sites were systematically introduced into a dCas9 gene with flanking BsaI sites using a targeted oligonucleotide library and recombineering (Fig. S1) ^37,38^. Second, these plasmid libraries were isolated, digested respectively with BsaI and either NheI or SpeI, and then ligated together (Fig. S1B). The resulting ligation of gene fragments produces deletions, as well as duplications, such that a MISER library has a triangular distribution, with near-wildtype (WT) length proteins most frequent and the largest deletions least frequent (Fig. 1C). To empirically determine the size range of functional deletions, the dCas9 MISER library was separated on an agarose gel and divided into six sublibrary slices of increasing deletion size. The sublibraries were then independently cloned into expression vectors and assayed for bacterial CRISPRi GFP repression via flow cytometry (Fig 1B, S2) ^39,40^. Sublibrary Slice 4 (ranging 3.2-3.5 kbp) was the most stringent (i.e. smallest) library with detectable repression, and functional variants became more frequent in slices possessing smaller deletions, as expected.

Fluorescence-activated Cell Sorting (FACS) and sequencing of MISER variants identified dCas9 deletion variants competent for DNA-binding. To focus sequencing on functional variants, Slice 4 and Slice 5 were sequenced pre- and post-FACS sorting, and the enrichment or depletion of individual variants was quantified (Fig S3). Four large deletion regions were independently identified in both libraries. Although the libraries target different size regimes, their overlapping data were significantly correlated (Fig S3). These data were normalized and combined to generate a comprehensive landscape of functional dCas9 deletions (Fig. 1C). 80% of sequencing depth was focused on deletions from 150 to 350 amino acids in length (Slice 4), and 51.4% (115,530/224,718) of these deletions were detected. Overall, this landscape includes data for 27.5% of all possible dCas9 deletions (257,737/936,396). The four large deletion regions roughly corresponded to the REC2, REC3, HNH, and RuvC-III domains. While larger deletions are bounded between domain termini, small deletions and insertions (~10 amino acids) are tolerated in much of the structure (Fig. S4), a finding that has been previously observed in other proteins ^17 22^. Two clear exceptions are the mechanistically essential ‘bridge helix’ ^35^, which orders and stabilizes the R-loop ^41,42^, and the ‘phosphate lock loop’ ^43^, which interacts with the PAM-proximal target strand phosphate to enable gRNA strand invasion. It should be noted that the enrichment data presented here is somewhat sparse and only a relative measurement of CRISPRi function; the larger-scale features of acceptable domain and sub-domain level deletions were therefore extensively validated with further *in vivo* and biochemical assays.

### Cas9 tolerates large domain deletions while retaining DNA-binding function

To validate the deletion profile, individual variants from each of the four deletion regions were either isolated from the library (Fig. S5) or constructed via PCR and assayed individually. Representative variants from each of the four deletion regions could be identified that exhibited bacterial CRISPRi nearly as effectively as full-length dCas9 (Fig. 2A, S6). Intriguingly, there are regions within our identified deletions that have been previously tested based on rational design, providing additional insight into the biochemical mechanisms lost with the removal of each domain ^35,44^. The most obvious of the acceptable deletions is of the HNH domain that is responsible for cleaving the target strand and gating cleavage by the RuvC domain; it is thus of little surprise that deletions of HNH are tolerated in a molecule that is required to bind but not cleave DNA. In fact Sternberg et al. previously demonstrated that a HNH-deleted (Δ768-919) Cas9 is competent for nearly WT levels of binding activity, but is unable to cleave ^45^. In contrast, we also uncovered a deletion in the RuvC-III domain that has never been observed. Modeling this deletion on the previously determined structure of SpCas9 bound to a DNA-target (PDB ID 5Y36) ^46^ revealed that it removes a large set of loops, an alpha helix and two antiparallel beta sheets (Fig S7). This deletion does not seem to overlay with a known functional domain and thus may serve as a module that further stabilizes the RuvC domain as a whole. Additionally, this deletion abuts the nontarget and target strand DNA (~4-6 Å) and may provide a highly useful site to replace with accessory fusions, such as deaminases suitable for base editing the non-target strand.

**Figure 2:**
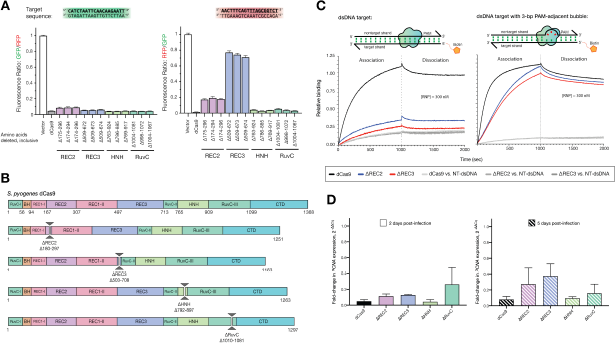
Cas9 tolerates whole-domain deletions while maintaining target-binding activity. **A)** *In vivo* transcription repression activity of MISER-dCas9 variants with specified amino-acids deleted, targeting either GFP (left) or RFP (right). dCas9s with REC2, REC3, HNH, or RuvC domain deletions have near-WT binding activity when targeted to GFP. When targeted to RFP, ΔREC3 and ΔREC3 show less robust binding activity. Data are normalized to vector-only control representing maximum fluorescence. Data are plotted as mean±SD from biological triplicates. **B)** Schema showing cloned MISER constructs with individual domain deletions corresponding to tolerated regions found in MISER screen. **C)** Bio-layer interferometry (BLI) assay of MISER constructs. ΔREC2 and ΔREC3 exhibit weak binding against a fully-complementary dsDNA target. Binding is rescued to near-WT levels, although at a slower rate, when the dsDNA contains a 3-bp bubble in the PAM-proximal seed region. Data are normalized to dCas9 binding to fully-complementary dsDNA. **D)** Measurement of CRISPRi efficacy of single-deletion MISER constructs in mammalian U-251 cells using RT-qPCR. U-251 cells were stably transduced with lentiviral vectors encoding dCas9 or MISER constructs fused with a KRAB repressor, along with lentivirus expressing sgRNA targeting *PCNA*. Cells were harvested 2 (left panel) or 5 (right) days post-transduction of the sgRNA and assayed for PCNA expression. Bar graphs represent fold-change of PCNA expression relative to a non-targeting sgRNA. Error bars represent SD for at least 2 replicates.

Our observations for the REC2 and REC3 domains likewise expand upon two rationally engineered deletions. Chen et al. previously demonstrated that the REC3 domain gates the rearrangement of the HNH cleavage by sensing the extended RNA:DNA duplex ^44^. Deletion of this domain (Δ497-713) ablated cleavage activity while maintaining full binding affinity. Nishimasu et. al. also previously deleted the REC2 domain because they postulated that it was unnecessary for DNA cleavage, as it is poorly conserved across other Cas9 sequences and lacks significant contact to the bound guide:target heteroduplex in the structure; however, the deletion mutant was found to have reduced activity ^35^.

To further validate the function and potential deficits of these single whole-domain deletions, we biochemically analyzed representative deletions of each of the REC2, REC3, HNH, and RuvC domains (Fig 2B). These single-deletion constructs are henceforth referred to as ΔREC2 (residues 180-297 deleted), ΔREC3 (Δ503-708), ΔHNH (Δ792-897), and ΔRuvC (Δ1010-1081) (Fig S9). Deletion variants were expressed, purified (see Supporting Information and Fig S9 for purification data), and assayed for DNA binding activity using bio-layer interferometry (BLI) (Fig 2C). Binding assays revealed that the REC2 deletion confers a defect in binding to a fully-complementary double-stranded DNA target (dsDNA) when complexed with a single-guide RNA (sgRNA). Interestingly, the defect is almost fully rescued upon the addition of a 3-bp mismatch bubble between the target and nontarget DNA strands adjacent to the PAM. DNA unwinding is initiated by Cas9 at the PAM-adjacent seed region, enabling the RNA-DNA R-loop hybrid to form. Rescue via seed bubble therefore suggests a potential role for the REC2 domain in unwinding dsDNA.

A similar phenomenon is observed with the ΔREC3 variant, although the binding defect is less pronounced than in ΔREC2. ΔREC3 is also unable to bind fully-complementary dsDNA - an effect that is rescued by the same PAM-adjacent 3-bp bubble in the dsDNA substrate, implying a similar DNA unwinding function by the REC3 domain. These results suggest that both the REC2 and REC3 domains are not essential for DNA binding by SpCas9, but may have evolved as “enhancer” domains to allow SpCas9 to more efficiently bind DNA inside the cell.

When measuring the repression activity of the ΔREC3 constructs *in vivo*, we also observed that the ΔREC3 appears to exhibit varying levels of repression between different gRNAs sequences. Specifically, we found that a GFP-targeting gRNA repressed stronger than an RFP-targeting gRNA with ΔREC3, after controlling for cell growth and fluorophore maturity (Fig. 2A). This was unexpected, since the binding of WT Cas9 is generally thought to be gRNA sequence-agnostic ^47^. One possibility is that the GC content of the targets in GFP and RFP could affect function, for example, a higher proportion of GC base-pairing in the “seed” region of a DNA target could present a greater energetic cost of unwinding to a deletion variant like ΔREC3^48^. Analysis of 16 additional spacer sequences and their repression activity relative to WT suggests this mechanism only moderately (R^2^ = 0.2) explains the variance (Fig S8). Further comprehensive analysis of the sequence-dependent variability is required to identify the precise energetic threshold the ΔREC3 construct overcomes to unwind DNA.

To test whether the MISER constructs retain DNA binding activity in mammalian systems, we performed CRISPRi to knock down genes in a U-251 glioblastoma cell line. We transduced target cells with lentiviral vectors expressing our single-deletion MISER constructs (ΔREC2, ΔREC3, ΔHNH, and ΔRuvC) fused to the KRAB repressor domain, followed by selection on puromycin. Stable cell lines were then transduced with a secondary lentiviral vector expressing an sgRNA targeting *PCNA*. After harvesting cells 2- and 5-days post-transduction of the sgRNA vector, we performed RT-qPCR to measure repression of the gene and assess efficacy of the MISER constructs in U-251 cells (Fig. S11). RT-qPCR of PCNA showed that 2 days post-transduction of the sgRNA the ΔREC2, ΔREC3, ΔHNH and ΔRuvC constructs repress PCNA expression relative to a non-targeting gRNA (sgNT) (Fig 2D; all measurements are averages ± S.D. from biological duplicates), with a mean fold-change of 0.11±0.03, 0.13±0.01, 0.04±0.03, and 0.26±0.21 in PCNA expression, respectively. At 5 days post-transfection, ΔREC2 and ΔREC3 appear to lose some repression activity (0.3±0.2 and 0.4±0.2 fold-change relative to sgNT, respectively), while the ΔHNH and ΔRuvC constructs are comparable to WT dCas9 at Day 5 (0.10±0.02 and 0.2±0.1 respectively) (see Supporting Information and Fig. S11 for more details on RT-qPCR).

### Stacking MISER deletions results in minimal DNA-binding CRISPR proteins

Protein domains are accreted during the evolution of large proteins ^3,4,49^. In principle, accretion could be experimentally reversed provided sufficient modularity is present to offset evolutionary divergence, epistasis, and other deleterious effects in ‘stacked’ deletions. To emulate this process while also engineering a minimal Cas9-derived DNA-binding protein, we generated a library of constructs that consolidated the ΔREC2, ΔREC3, ΔHNH, and ΔRuvC deletions found by the MISER screen.

A library of multi-deletion variants, termed CRISPR Effectors (CE) due to their highly pared-down sequence relative to wild-type Cas9, were constructed as follows: individual sublibraries of deletions from REC2, REC3, and the HNH domains were isolated from the full MISER library. This was done by selecting against the full-length dCas9 sequence by targeting a pre-existing restriction site within each deleted region, so that only transformations of circular plasmids that had the respective deletion would be favored (Fig. 3B, S5). The RuvC deletion was an exception since it did not have a pre-existing restriction site; therefore a manually constructed ΔRuvC variant (Δ1010-1081) was amplified and used as a starting point for further stacking.

**Figure 3:**
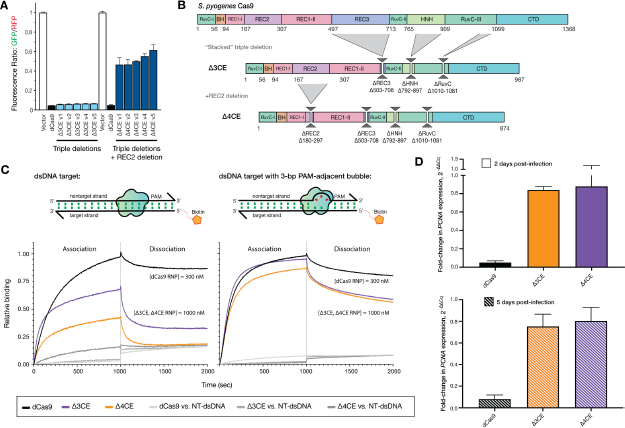
Stacking multiple domain deletions on Cas9 results in defective DNA-binding activity. **A)** *In vivo* transcription repression activity of MISER CRISPR effectors containing triple (Δ3CE) and quadruple (Δ4CE) deletion variants. Sublibraries of REC2, REC3, HNH, and RuvC were combined to build a library of stacked deletions, and the resulting library was assayed for high-performing variants using FACS (light blue bars). As none of the variants contained a REC2 deletion (~Δ167-307), we named the highest-performing triple-deletion variant in this library (Library 2; see Fig S6) Δ3CE. To force a library containing REC2 deletions, a sublibrary of REC2 deletions was added to Δ3CE, resulting in a library of quadruple deletion variants that contain Δ3CE and a REC2 deletion (dark blue bars). Data are plotted as mean±SD from biological triplicates. **B)** Expression constructs for Δ3CE and Δ4CE, with specified deletions manually cloned in. **C)** BLI assay of CE constructs. Δ3CE and Δ4CE exhibit almost no binding against a fully-complementary dsDNA target at 300 nM RNP (see Fig. S10); and weak binding at 1000 nM RNP. Binding is rescued to near-WT levels when RNP concentration is 3.3x that of dCas9 if the dsDNA contains a 3-bp bubble in the PAM-proximal seed region. Data are normalized to 300 nM dCas9 binding to fully-complementary dsDNA. **D)** Measurement of CRISPRi efficacy of Δ3CE and Δ4CE in U-251 cells using RT-qPCR. Fold-change in PCNA expression levels is measured by RT-qPCR, 2- and 5-days after KRAB-Δ3CE and -Δ4CE expressing cell lines are transfected with a sgRNA targeting *PCNA*. Δ3CE and Δ4CE exhibit weak DNA binding and transcriptional repression activity compared to dCas9. Bars represent fold-change of PCNA expression relative to a non-targeting sgRNA. Error bars represent SD for at least 2 replicates.

The dCas9 gene was divided into four fragments spanning the major deletions and recombined using Golden Gate cloning (Fig. S6). The resulting library, CE Library 1, was assayed using bacterial CRISPRi, and functional variants were isolated by FACS, as above. A variety of functional CEs were obtained (Fig. 3A), although surprisingly, none of them possessed a REC2 deletion. We therefore generated a second library, CE Library 2, in which a library of triple-deletion variants were cross-bred with REC2 deletion variants to ensure diversity in deletions from this region (Fig. S6). Again, the most functional CE variants isolated by FACS did not contain REC2 deletions. Finally, in an attempt to force a minimal CE, the most active CE variant from CE Library 1 and 2, termed Δ3CE, was directly combined with a library of REC2 deletions and screened for activity. The resulting ‘hard-coded’ quadruple deletion CE variants all exhibited loss of function (Fig. 3A), which explains why the REC2 deletion was lost in our functional variants. The most active variant (Δ4CE) possessed a deletion of Δ180-297 and was confirmed upon re-transformation to display ~50% the activity of WT dCas9 (Fig. 3A, 3C) in *E. coli*.

To validate the stacked deletion constructs biochemically, we expressed and purified the Δ3CE and Δ4CE variants from *E. coli* (Fig. 3B, S10). BLI experiments revealed that compared to the bacterial *in vivo* repression data, the DNA-binding abilities of both stacked deletion constructs were attenuated relative to dCas9 (Fig 3C). To obtain reasonable kinetic profiles, the concentration of RNP for Δ3CE and Δ4CE was increased to 1000 nM, but even under these conditions both variants lag WT dCas9 at 300 nM. The PAM-interrogation ability of the two constructs appeared to be intact, as evidenced by the sharp drop-off in signal during the dissociation phase, but both Δ3CE and Δ4CE dissociated at a much higher rate compared to dCas9. The k_on_ was restored upon addition of a 3-bp bubble, suggesting that these minimal Cas9s possess the kinetic defect in dsDNA binding inherent to both ΔREC2 and ΔREC3. The fact that these minimal constructs are still able to bind DNA in a sgRNA-targeted fashion is surprising, considering that the Δ3CE and Δ4CE constructs retain only ~72% and ~63%, respectively, of the original dCas9 protein primary sequence (Fig. 3B).

We assessed the DNA binding activity of the CE constructs in mammalian cells similarly to the single-deletion variants described earlier. As before, we performed CRISPRi against PCNA in U-251 cells, this time transducing the Δ3CE and Δ4CE KRAB fusions and sgRNA, followed by harvesting and RT-qPCR 2- and 5-days post-transfection. Unlike the single-deletion variants, Δ3CE and Δ4CE do not repress nearly as well as dCas9, exhibiting a fold-change in PCNA expression relative to non-targeting sgRNA of 0.75±0.11 and 0.80±0.13, respectively, after five days post-transduction of the sgRNA (Fig. 3D). This result suggests that the Δ3CE and Δ4CE constructs are functional but severely defective in DNA binding in a mammalian system.

### The minimal Δ4CE construct has a similar structure as an ablated wild-type SpCas9

To understand the structural rearrangement accompanying domain deletion, we used single-particle cryo-EM to determine a reconstruction of the Δ4CE DNA-bound holocomplex (Fig. 4), to a resolution of 6.2 Å (Fig. S12). Remarkably, overlaying the density of the Δ4CE construct over the WT SpCas9 R-loop structure (PDB ID 5Y36) ^43^ as a rigid-body model shows that the minimal complex, consisting primarily of the REC1, RuvC, and C-terminal domains, possesses the same overall architecture as the WT holocomplex (Fig 4A, S12). The double-helical dsDNA target and the stem-loop of the gRNA that are part of the R-loop can be resolved from the density and overlays almost exactly over the WT SpCas9 R-loop. This observation supports the hypothesis that the R-loop is a thermodynamically stable structure that drives the formation of the primed Cas9 RNP-DNA complex ^50,51^. Although individual residues cannot be resolved, the remaining RuvC domain in the construct is linked to the C-terminus of the REC1 domain via a TS linker (MISER scar), thereby maintaining a bi-lobed complex reminiscent of WT SpCas9. The gRNA-interacting regions of the REC1 and CTD are also spatially conserved, consistent with their observed indispensability on the MISER enrichment map. This raises the question of how the minimal protein is able to form a stable R-loop despite lacking a large part of the REC lobe.

**Figure 4:**
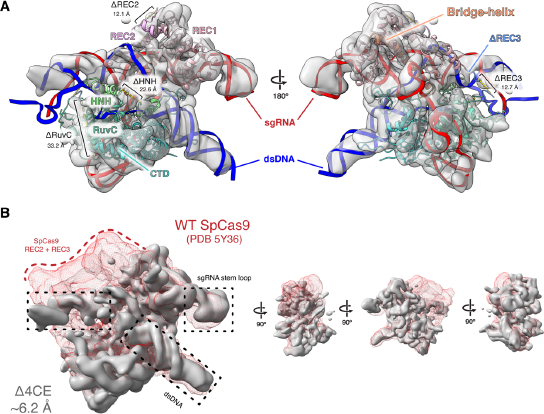
Density map of Δ4CE compared to WT SpCas9. **A)** Single-particle cryo-electron microscopy was used to obtain a density map of the dsDNA-bound RNP complex of the Δ4CE construct at an overall resolution of 6.2 Å (EMD-22518). Light grey volume shows the Δ4CE density overlaid onto RNA-DNA hybrid R-loop (red and blue) and structure of WT SpCas9 (PDB 5Y36). Cartoon model corresponds to the WT SpCas9 structure, showing only the remaining residues and corresponding domains after the REC2, REC3, HNH, and RuvC deletions from the Δ4CE construct are manually removed from the model. Deletion termini are labeled with the distances between termini. **B)** Density of Δ4CE cryo-EM overlaid with the WT SpCas9 clearly shows volumes representing dsDNA target and the sgRNA stem-loop (black boxes). The red mesh represents the total WT SpCas9 density from EMD-8236.

## Discussion

Protein evolution takes large steps through sequence space using domain rearrangements, duplications, and indels ^2,52^. While rearrangements, duplications, and insertions have been widely studied, domain deletions are largely under-investigated, due to limited experimental data and the difficulty in properly annotating deletions in protein sequence datasets^53^. Although deletion studies in proteins have been performed, they are limited in their scope in regard to the scale of deletions, complexity, and generalizability. In this work, we present a technique that is versatile, comprehensive, and unbiased to probe the deletion landscape of virtually any protein, limited only by the fidelity and efficiency of a functional screen.

We have used SpCas9 as proof-of-concept to demonstrate the utility of MISER because it is a well-characterized, multi-domain protein, easy to assay, and its overall size poses a limit for therapeutic delivery. The wild-type SpCas9 gene is too large to be packaged into an adeno-associated viral vector (AAV), which has a maximum reported cargo size of <5 kb^54,55^ when including the sgRNA sequence and necessary promoters. There are now smaller characterized CRISPR-Cas effectors suitable for AAV delivery by themselves ^19,56^, but an important need in both research and therapy is delivery of effectors fused to other domains, such as for base-editing and transcriptional activation or repression ^57^. MISER may thus find utility in minimizing these much larger constructs. Additionally, immunogenicity is emerging as a major issue when developing SpCas9 as a therapeutic, and deleting antigenic surface residues can potentially reduce the reactivity of the protein against the immune system ^58,59^.

We were surprised to discover the effect the deletion of the REC2 domain had on SpCas9 binding. Nishimasu et al. had previously reported that a REC2 deletion (Δ175-307) retained ~50% of editing activity, and suggested that the attenuated activity might be due to poor expression or stability ^35^. In contrast, our data suggest that the ΔREC2 variant folds and retains target recognition and binding function but loses DNA unwinding capability. The observation that ΔREC2 binding is restored upon addition of a 3-bp bubble adjacent to the PAM suggests that the poor binding is due to a kinetic defect. The specific nature of the defect requires further study, although we speculate that the REC2 domain interacts nonspecifically and transiently with the R-loop, perhaps stabilizing the DNA strands during hybridization (i.e. lowering the kinetic barrier) or stabilizing the final R-loop complex (i.e., lowering the energetic cost of unwinding and hybridization) ^44^.

We also note the observed difference in activity of the MISER constructs between bacterial *in vivo* repression experiments and the *in vitro* binding activity using BLI. We speculate that the MISER constructs are inherently defective for binding target DNA, but that sufficiently perturbed dsDNA in bacteria—such as during replication, transcription, or other rearrangements—presents enough opportunity in the form of dynamically un- and under-wound dsDNA, or stretches of single-stranded DNA, to allow the gRNA to anneal to the spacer sequence ^51,60^.

Finally, in our cryo-EM structure of Δ4CE, we note the remarkable similarity of the protein to WT SpCas9, which underscores the inherent stability of the Cas9 R-loop complex. Previous studies have shown that formation and maintenance of the R-loop is the molecular “glue” that holds the DNA-RNA-protein complex together ^50^. The similitude between the WT and Δ4CE structure also hints at the evolutionary history of SpCas9, suggesting that the “essential” function of the protein was to enable the formation of an R-loop upon a RuvC scaffold for DNA binding and cleavage, which was then tuned by accretion and interactions of other domains—such as those that comprise the REC lobe and the HNH domains ^9,61^. Notably, this analysis ignores the role of the gRNA; future iterations of MISER could also be used to evaluate the deletional landscape of CRISPR-associated RNAs.

MISER facilitates the study of protein deletions with unprecedented versatility and efficiency. In this study we have explored domain modularity and essentiality of CRISPR-Cas9 domains, but MISER can be adapted to any application requiring a reduction in genetic size. AAV-based transgene delivery is subject to a < 5 kb payload limit and is a prime target for MISER. Besides CRISPR proteins and their cognate gRNAs, there are numerous other therapeutic proteins limited by their size, such as CFTR (cystic fibrosis) and dystrophin (muscular dystrophy)^62 54^. Beyond threshold effects, even partially reducing the size of AAV genomes can provide a large advantage in packaging efficiency by improving capsid formation^55^. Finally, MISER also reveals small deletions tolerated within proteins, which suggests that this approach could be useful in the development of non-immunogenic biomolecules. Paring away antigenic residues may remove antigenic epitopes on a protein surface, thus allowing the molecule to function without eliciting an immune response ^63,64^.

## Supporting information

Auxiliary Supplemental Materials

## Acknowledgments

This work was supported by NIH grants 1R01GM127463 (D.F.S.) and RM1HG009490 (J.A.D., D.F.S). Additional support and reagents were provided by Agilent Technologies. A.S. was supported by the NSF GRFP (grant no. 1752814), S.A.H. was supported by NIH Training Grant 5T32GM066698-10, and M.A. was supported by the ARCS Foundation. C.F. was supported by a U.S. NIH K99/R00 Pathway to Independence Award (K99GM118909, R00GM118909) from the NIGMS. J.A.D. is an Investigator of the Howard Hughes Medical Institute (HHMI), and this study was supported in part by HHMI. This work used the Vincent J. Coates Genomics Sequencing Laboratory at University of California, Berkeley, supported by NIH S10 instrumentation grants (S10RR029668 and S10RR027303). We thank Mary West and the CIRM/QB3 Shared Stem Cell Facility/High-Throughput Screening Facility for technical support, as well as Timothy Brown (Thermo Fisher Scientific) for flow cytometry support. We thank Daniel Toso and Paul Tobias for assistance with cryo-EM data collection at Bay Area Cryo-EM facility at the University of California, Berkeley. We also thank Emeric Charles, Shin Kim, Rob Nichols, Luke Oltrogge, Avi Flamholz, and Joshua Cofsky for technical support and productive discussions.

## Data Availability Statement

All custom scripts are available at https://github.com/savagelab. All sequencing data that support the findings of this study are available from the authors upon reasonable request. Cryo-EM data are available at the Electron Microscopy DataBank (EMDB); accession code 22518. All other relevant data are available from the corresponding author on request.

