## Supporting Information for "Comprehensive deletion landscape of CRISPR-Cas9 identifies minimal RNA-guided DNA-binding modules"

Affiliations:
[1]Department of Molecular and Cell Biology, University of California, Berkeley, Berkeley, CA 94720, USA
[2]Innovative Genomics Institute, University of California, Berkeley, Berkeley, CA 94720, USA
[3]Scribe Therapeutics, Alameda, CA 94501, USA
[4]Gladstone Institutes, San Francisco, CA 94158, USA
[5]Department of Cellular and Molecular Pharmacology, University of California, San Francisco, San Francisco, CA 94158, USA
[6]Division of Biological Sciences, University of San Diego, San Diego, CA 92093, USA
[7]Graduate Group in Biophysics, University of California, Berkeley, Berkeley, CA 94720, USA
[8]Department of Bioengineering, University of California, Berkeley, Berkeley, CA 94720, USA
[9]Howard Hughes Medical Institute, University of California, Berkeley, Berkeley, CA 94720, USA
[10]Molecular Biophysics and Integrated Bioimaging Division, Lawrence Berkeley National Laboratory, Berkeley, CA 94720, USA
[11]Department of Chemistry, University of California, Berkeley, Berkeley, CA 94720, USA

* These authors contributed equally to this work.

### Experimental Design

#### Molecular Biology

All restriction enzymes were ordered from New England Biolabs (NEB). Polymerase Chain Reaction (PCR) was performed using Q5 High-Fidelity DNA Polymerase from NEB. Ligation was performed using T4 DNA Ligase from NEB. Agarose gel extraction was performed using the Zymoclean Gel DNA Recovery kit, and PCR clean-up was performed using the 'DNA Clean & Concentrator', both from Zymo Research. Plasmids were isolated using the QIAprep Spin Miniprep Kit (Qiagen). All DNA-modifying procedures were performed according to the manufacturers' instructions.

#### MISER library construction: Plasmid Recombineering

Two sets of 1368 oligonucleotides were designed and ordered as Oligonucleotide Library Synthesis (OLS) from Agilent Technologies (Table S1). Oligonucleotides were designed to insert a six base pair (bp) recognition sequence for either the restriction enzyme NheI or SpeI between every codon in dCas9 (Figure S1A). The full list of ordered oligonucleotides is available as Auxiliary Supplementary Materials - Recombineering Oligonucleotides. Internal priming sites were included in order to amplify NheI or SpeI specific oligonucleotide libraries. A modified amplification procedure was performed. In a 50 µL PCR reaction, 10 ng of template oligonucleotide library was amplified according to manufacturer's instructions, but with an extension time of only five seconds, and a total of only 15 cycles. 1.5% dimethyl sulfoxide (DMSO) was also included in the PCR reaction. These modifications were empirically determined in order to minimize undesirable higher order PCR products that were observed to be produced by amplification. These side products are likely the result of complementary oligonucleotides priming one another. Notably this phenomenon is likely inherent to amplification of a library of DNA tiled across a common sequence--in this case dCas9. PCR primers can be found in Table S6 and Auxiliary Supplementary Materials – Primer Sequences. 24 such reactions were typically performed in parallel and then combined, followed by concentration with Zymo DNA Clean & Concentrator. BsmbI restriction digestion was then used to remove priming ends, followed by a second concentration with Zymo DNA Clean & Concentrator, resulting in mature double-stranded recombineering-competent DNA.

Plasmid recombineering was performed as described in Higgins et al. 2017, using strain EcNR2 (Addgene ID: 26931) to generate MISER libraries in plasmid pSAH060. Plasmid sequences can be found in Auxiliary Supplementary Materials – Plasmid Sequences. Briefly, mature double-stranded recombineering-competent DNA at a final volume of 50 µL of 1 µM, plus 10 ng of pSAH060, was electroporated into 1 mL of induced and washed EcNR2 using a 1 mm electroporation cuvette (BioRad GenePulser). A Harvard Apparatus ECM 630 Electroporation System was used with settings 1800 kV, 200 Ω, 25 µF. Three replicate electroporations were performed, then individually allowed to recover at 30° C for 1 hr in 1 mL of SOC (Teknova) without antibiotic. LB (Teknova) and kanamycin (Fisher) at 60 µg/mL was then added to 6 mL final volume and grown overnight. A sample of recovered culture was diluted and plated on kanamycin to estimate the total number of transformants, typically >$10^7$. Cultures were miniprepped and combined the next day. Plasmid recombineering is relatively inefficient, and only a fraction of recovered plasmids contained successful NheI or SpeI insertions. In order to recover completely penetrant libraries, an intermediate cloning step was performed. A PCR product conferring resistance to chloramphenicol was cloned into both libraries of pSAH060 plasmids (Auxiliary Supplementary Materials - Chloramphenicol Selection). This PCR product contained either flanking

NheI restriction sites or SpeI restriction sites, such that only modified pSAH060 plasmids (possessing NheI or SpeI restriction sites) could obtain chloramphenicol resistance through NheI/SpeI digestion and subsequent ligation. Libraries were then purified (Zymo) and transformed into XLI-Blue competent cells for overnight selection in chloramphenicol (Amresco) at 25 µg/mL, followed by plasmid isolation the next day. Samples of recovered cultures were also plated on both kanamycin alone (native pSAH060 resistance) and chloramphenicol alone (resistance mediated by successful recombineering insertion) to estimate the fraction of modified plasmids and therefore the restriction library size. Recombineering efficiencies were observed at ~0.5% by this method, indicating restriction library sizes of ~50,000, well above the number of unique insertion sites per library (1,368). Finally, chloramphenicol resistant pSAH060 libraries were digested with either NheI or SpeI as appropriate, removing the chloramphenicol cassette. The libraries were run on an agarose gel, and the 5953 bp (5947 bp pSAH060 + 6 bp inserted restriction site) linear band corresponding to each library was gel extracted. To construct deletion variants composed of N- and C- terminal dCas9 fragments, one µg of each library was mixed and digested with BsaI, then cleaned up (Zymo). The resulting DNA mixture contained equimolar free dCas9 N- and C-terminal fragments, as well as equimolar pSAH060 vector backbone. This mixture was then ligated in the presence of SpeI and NheI, 'locking' dCas9 fragments together by one of two six bp scar sites not recognized by either enzyme (Figure S1B). The ligated MISER library was transformed into XL1-Blue, grown overnight and plasmids were isolated the next day. The MISER library of dCas9 is quite large, with 936,396 possible deletions (N(N + 1) / 2, N = 1368), and all cloning steps were performed with validation that >$10^7$ transformants were obtained.

**MISER library construction: library size selection**

The MISER library is theoretically composed of all possible N- and C-terminal fragments, including both duplications and deletions. To isolate deletions in a particular size range, the MISER library was digested with BsaI, in order to excise the dCas9 gene from the vector backbone, and run on an agarose gel. Various slices of the MISER library were individually gel extracted (Fig. S2A), ligated into expression vector pSAH063 (Fig. S2B), and transformed into *E. coli*.

**Fluorescence repression assays and flow cytometry**

The catalytically dead dCas9 MISER variants were used to repress the transcription of genomically encoded fluorescent reporter genes in *E. coli* as previously described [1]. A sgRNA targeting Green Fluorescent Protein (GFP) was transcribed from plasmid pgRNA-bacteria (Addgene ID 44251) [1], which results in repression of constitutively expressed GFP, contingent on functional dCas9 expression from pSAH063 [2]. This repression was quantified relative to non-targeted Red Fluorescent Protein (RFP), which is expressed from the same genomic locus [1]. This assay yields robust repression detection (Fig. S2B), with at least an order of magnitude lower GFP signal after 8 hours of growth at 37° C with 750 rpm shaking in LB media + 1 nM Isopropyl β-D-1-thiogalactopyranoside (IPTG) induction of dCas9 from pSAH063. Assays and flow cytometry were conducted in either an M1000 plate reader (Tecan) or an SH800 Cell Sorter (Sony Biotechnology). For GFP/RFP ratiometric measurements (Fig. 2A, 3A) there was no significant difference between samples for the RFP fluorescence measurement.

#### Deep sequencing

100 nucleotide single end reads were used to sequence the dCas9 Slice 4 and Slice 5 libraries. dCas9 open reading frames were amplified from pSAH064 libraries with primers SAH_356 and SAH_358. PCR products were further prepared for deep sequencing by the UC Berkeley Functional Genomics Laboratory. Sequencing was performed by the UC Berkeley Vincent J. Coates Genomics Sequencing Laboratory on an Illumina HiSeq4000. Samples were mixed at custom ratios as follows: Slice 5 Naïve Library – 10%; Slice 5 Sorted Library – 10%; Slice 4 Naïve Library – 40%; Slice 4 Sorted Library – 40%. Sequencing analysis was performed with custom MATLAB scripts available online at https://github.com/savagelab. Briefly, reads were analyzed for the novel presence of the two possible MISER scar sequences, 'GCTAGT' or 'ACTAGC'. The majority of reads were fully WT dCas9 sequences, as expected due to the fact that scar sequences can occur anywhere along dCas9. Once detected, reads containing 15 bp upstream and downstream of the scar (that exactly matched dCas9 sequence) were used to identify the location of a deletion. Sequencing statistics can be found in Table S3. Enrichment ratios were calculated by taking the ratio of the frequency of each variant before and after selection [3]. To conservatively display variants only detected in one library, one artificial read was added to both datasets. The log base ten of these enrichment ratios were plotted (Figure S3 A and B) for each of the two libraries. For visualization, these two datasets were also normalized according to their Pearson Correlation (Figure S3 E), combined (the mean was calculated for those variants with two values), and rescaled for display (Figure 1C and S4 A).

#### Protein expression and purification

A *Streptococcus pyogenes* Cas9 gene containing nuclease-deactivating mutations D10A/H840A (a.k.a. dCas9) was cloned into a pET14b expression vector, encoding a N-terminal 6xHis fusion tag and a C-terminal 2xNLS fusion tag. Specific MISER dCas9 variants were cloned by PCR-amplification (Q5 High-fidelity polymerase, NEB) of the dCas9 gene excluding deleted regions obtained from MISER screen (see Table S4 for primer sequences). Plasmids were verified by Sanger sequencing (UC Berkeley DNA Sequencing Facility). dCas9 and MISER constructs were overexpressed in *E. coli* BL21 (DE3) LOBSTR expression system (Kerafast). Cells were grown in Terrific Broth, modified media with 8 mM $MgCl_2$ and 0.5 glycerol and induced at ~0.6 OD with 0.5 mM IPTG. Cells were resuspended in Lysis Buffer (20 mM HEPES pH 7.5, 1 M KCl, 15 mM imidazole, 1 mM TCEP, 10% glycerol, 0.1 mM PMSF, Roche protease inhibitor tablet), lysed by sonication and clarified by centrifugation, and incubated with Ni-NTA resin to purify soluble fractions. Protein-bound Ni-NTA resin was washed with Wash Buffer (Lysis Buffer + 0.1% Triton X-114), and eluted (Elution Buffer: 20 mM HEPES pH 7.5, 150 mM KCl, 300 mM imidazole, 1 mM TCEP, 10% glycerol). Eluted fractions were subjected to a Heparin Sepharose column (GE Healthcare) for ion-exchange chromatography (300 mM KCl to 1 M gradient), concentrated, and further purified on a gel-filtration column (Superose 6 Increase, GE Healthcare). Protein Storage Buffer was as follows: 20 mM HEPES pH 7.5, 150 mM KCl, 1 mM TCEP, 10% glycerol. Purified protein aliquots were flash-frozen in liquid nitrogen and stored at -80°C. Concentrations were measured via Nanodrop A280 (ThermoFisher Scientific).

#### *In vitro* DNA binding assays

Purified proteins were complexed with 1.2x molar ratio sgRNA in the presence of 5 mM $MgCl_2$. 5'-biotinylated target DNA and corresponding non-target DNA was purchased from IDT as single-stranded oligos and annealed 1:1 according to standard IDT protocols. All bio-layer interferometry (BLI) measurements were performed on an Octet RED384 system (ForteBio). Biosensors coated with streptavidin (SA) were incubated in BLI Buffer (20 mM HEPES pH 7.5, 100 mM KCl, 5 mM $MgCl_2$, 10 µg/mL Heparin, 50 µg/mL bovine serum albumin, 0.01% v/v IGEPAL CA-630, 1 mM TCEP, 10% v/v glycerol) for ~10 min prior to assay. 5'-biotinylated target DNA (ligand) and corresponding non-target DNA was purchased from IDT as single-stranded oligos and annealed 1:1 according to standard IDT protocol (See Table S4 for oligo sequences).

Biotinylated dsDNA was diluted in BLI buffer to a concentration of 10 nM. dCas9 or MISER construct RNPs were diluted in BLI Buffer at various concentrations (0.1x to 10x reported $K_D$). BLI step sequence was as follows: SA biosensors were incubated in BLI buffer for 60 seconds (baseline); dsDNA ligands were loaded onto SA biosensors for 300 seconds (loading); SA biosensors were incubated in BLI buffer for 60 seconds again to re-equilibrate ligand-bound tip (baseline); dsDNA-functionalized biosensors were incubated with RNP analytes for 1000 seconds (association); and biosensors were incubated in baseline wells from Step 1 for 1000 seconds (dissociation). All steps were performed at 37° C with stirring (1000 RPM). Data analysis was performed with Octet Data Analysis HT software (ForteBio).

#### Mammalian CRISPR interference (CRISPRi) assay

For the mammalian CRISPR interference (CRISPRi) based competitive proliferation assay, human U-251 glioblastoma cells were stably transduced with lentiviral vectors (pSC066) expressing MISER or WT-dCas9 KRAB fusion proteins, followed by selection on puromycin (InvivoGen, #ant-pr-1; 1.0-2.0 µg/ml). The respective cell lines were then transduced with a secondary lentiviral vector (pCF221) expressing mCherry fluorescence protein and either CRISPRi sgRNAs targeting essential genes (sgPCNA, sgRPA1) or non-targeting controls (sgNT). After mixing with the respective parental population (at approximately an 80:20 ratio of transduced to non-transduced cells), the percentage of mCherry positive cells was monitored by flow cytometry (Attune NxT flow cytometer, Thermo Fisher Scientific) over several days to assess the effect of CRISPRi with the given Cas9-variant on cell proliferation. CRISPR interference (CRISPRi) sgRNAs had been previously designed [4], as were non-targeting sgRNAs [5]. The sgRNAs were designed with a G preceding the 20-nucleotide guide for better expression from U6 promoters and cloned into the pCF221 lentiviral vector for expression [6].

#### Reverse-transcription quantitative PCR (RT-qPCR)

To measure the efficacy of CRISPRi repression of essential genes by dCas9-MISER constructs in cultured mammalian cells, we performed RT-qPCR of targeted genes in human U-251 glioblastoma cells. Cells were stably transduced with lentiviral vectors encoding dCas9- or MISER-KRAB proteins, and sgRNA targeting PCNA (sgPCNA-i6) as described in the mammalian CRISPRi experiment (including non-targeting guide sgNT-1), except without any mixing with the parental population. Cells were allowed to grow and then harvested 2 and 5 days post-transduction. RNA was extracted using Trizol-chloroform and stored in -80° C [7]. RNA was reverse-transcribed to cDNA with RNA-to-cDNA EcoDry™ Premix with random hexamers (Takara Bio), using manufacturer's protocols. Quantitative PCR (qPCR) amplification of cDNA was performed using primers specific for *PCNA* (oAS089-92, Table S4) using SYBR Green PCR Master Mix (ThermoFisher Scientific) in a QuantStudio 3 Real-time PCR System (ThermoFisher Scientific). GAPDH was used as the housekeeping control (amplified with primers oAS117-118, Table S4). All results are reported relative to the expression of PCNA in cells transfected with non-target gRNA (sgNT-1, Table S4). Only amplification plots below a ΔRn threshold of 0.040 and a $C_t$ value <35 cycles were used for analysis of expression levels. $\Delta C_q$ values were calculated by subtracting Cq values of GAPDH amplifications from PCNA, and $\Delta\Delta C_q$ values were calculated by subtracting the non-target samples from the target samples. Fold-change in expression is reported as $2^{-\Delta\Delta Cq}$.

**Cryo-electron microscopy sample preparation and image acquisition**

The ternary complex was prepared at 37 °C using a Δ4CE, sgRNA, and dsDNA target at a ratio of 1:1.5:2 in complexing buffer (30 mM Tris-HCl, pH 8.0, 150 KCl, 5 mM $MgCl_2$, 5 mM DTT, 2.5 % glycerol). Protein and sgRNA were incubated for 30 minutes prior to addition of dsDNA for an additional 1 hour of incubation. The sample was then desalted using a spin-column (Zeba) into Complexing Buffer containing 0.1% glycerol to be used for grid preparation. To prepare the sample for imaging, 3.2 µL of the ternary complex (around 30 nM) was applied to R1.2/1.3 Cu 200 grids (Quantifoil) coated with a thin layer of homemade continuous carbon that had been glow-discharged for 15 s immediately before use. The sample was incubated on the grid at 100% humidity and 16 °C for 10 s prior to blotting for 5 s with filter paper and plunging into liquid ethane cooled to liquid nitrogen temperatures using a Vitrobot Mark IV (TFS). The sample was imaged using a Talos Arctica transmission electron microscope (TFS) operated at 200 kV and equipped with a K3 direct electron detector (Gatan) at the Bay Area Cryo-EM facility at the University of California, Berkeley. Movies were recorded in super-resolution counting mode at an effective pixel size of 0.45 Å, with a cumulative exposure of 60 $e^{-}\cdot Å^{-2}$ distributed uniformly over 60 frames. Automated data acquisition was performed using image-shift and active beam tilt compensation as implemented in SerialEM-v3.7 to acquire movies from a 3x3 array of holes per stage movement [8]. In total, 3400 movies were acquired with a realized defocus range of -1.5 to -3.8 µm.

**Cryo-EM image processing**

All steps were performed using RELION-v3.1b unless otherwise indicated [9]. Movies were motion-corrected, exposure-filtered, and Fourier cropped to a pixel size of 0.9 Å using and the initial CTF parameters estimated by CTFFIND-v4.1.13 [10]. Micrographs were culled by thresholding for CTF-fit resolutions better than 8 Å and manual curation to yield a set of 2554 micrographs used in further processing. An initial set of 97,827 particles were picked using the general model of Boxnet2 [11]. These particles were extract in a 256 pixel box Fourier cropped to 64 pixels (3.6 $Å\cdot px^{-1}$). Iterative rounds of reference-free 2D classification resulted in 85,327 particles, which were used to generate an ab initio 3D-reference by stochastic gradient descent. Particles were re-extracted and upsampled in a 128 pixel box (1.8 $Å\cdot px^{-1}$) for further processing. Unsupervised 3D classification did not resolve distinguishable classes. Thus, all particles were subjected to 'gold-standard' 3D auto-refinement using a reference low-pass filtered to 25 Å and a soft shape-mask. This yielded a reconstruction at a nominal resolution of 6.4 Å based on the FSC0.143 criterion and using phase-randomization to correct for masking artifacts [12]. This set of particles was then used to train a picking model with Topaz-v0.2.3 [13]. This approach resulted in a set of 288,416 particle coordinates. The new set of particles was extracted in a

128 pixel box (1.8 Å·px$^{-1}$) and subjected to reference-free 2D classification, which resulted in a set 167,245 particles. Additional attempts at 3D classification did not resolve distinguishable classes. This final set of particles was used for 3D auto-refinement as described above and resulted in a 6.2 Å reconstruction. Further processing using reference-based fitting of particle motion and CTF parameters did not yield improvements. Resolution anisotropy of the final reconstruction was assessed using the 3DFSC web server [14].

**Modelling of the cryo-EM map**

The previously published coordinate model for the 5.2Å cryo-EM structure of SpCas9 ternary complex (PDB ID 5Y36) was used as an initial model [15]. To this end, the protein domains were deleted from 5Y36 to match those of Δ4CE. The edited coordinate model was then docked as a rigid-body into the RELION post-processed map using ChimeraX-v1.0, which resulted in a cross-correlation value of 0.73 against a 6.2 Å map simulated from the coordinate model [16]. For display purposes, a denoised version of the Δ4CE map was generated with LAFTER as part of the CCPEM-v1.4.1 suite [17].

### Supporting Figures

**Figure S1.**

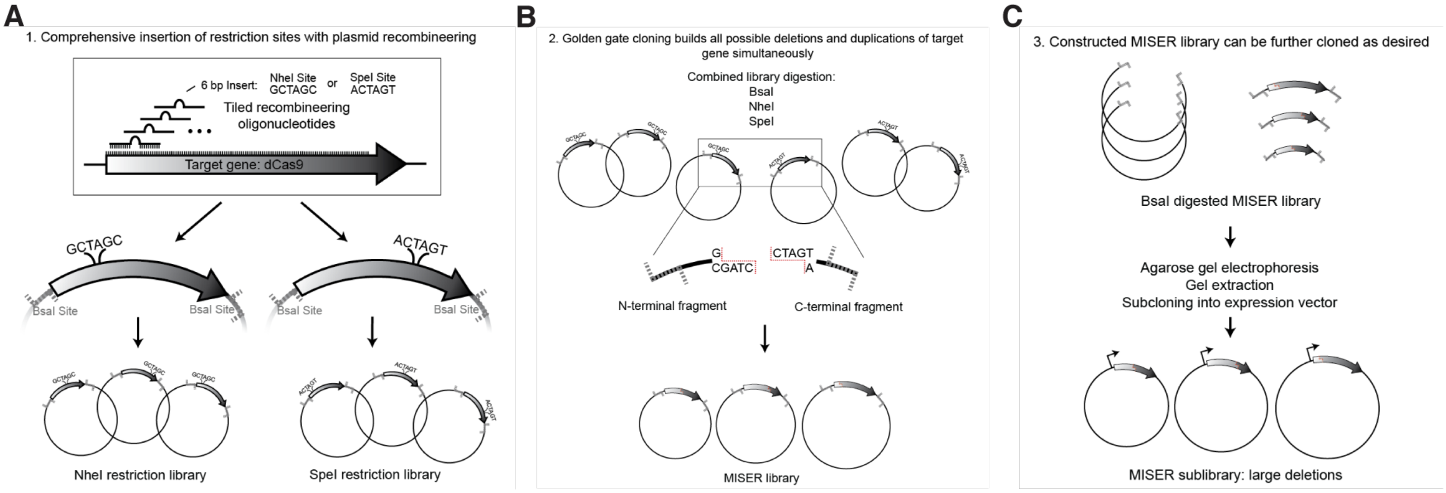

**<u>Figure S1:</u> Full cloning scheme for Minimization by Iterative Size-Exclusion and Recombination (MISER).** The method can be considered in three parts. **A)** Plasmid recombineering generates two comprehensive libraries of restriction site insertions across the target gene. These restriction sites are both novel to the target plasmid and produce compatible sticky ends. Recombineering was performed similarly as in (Higgins 2017), where the target gene lacks a promoter and start codon to prevent growth biases during library construction and is flanked by BsaI sites for later Golden Gate cloning (here, plasmid pSAH060). Additionally, rather than mutagenic oligos, double stranded PCR product was used for recombineering, and another cloning step was introduced to remove unmodified plasmids. These modifications are described in Experimental Design. **B)** Modified golden gate cloning generates a library of ligated N- and C- terminal fragments of the target gene, comprehensively producing protein deletion variants as well as duplication variants. An equimolar mixture of the two plasmid libraries is mixed and fully digested to produce free N- and C- terminal fragments of the target gene. This fragment mixture is then re- ligated in the presence of NheI and SpeI. Successful ligation of an N- and C-terminal fragment from differing libraries produces one of two possible 6 base-pair scar sequences. These novel scar sequences are not recognized by either NheI or SpeI, thus trapping the desired chimeric product as a final ligated vector. Because N- and C-terminal fragments are ligated randomly, these chimeric products produce both protein deletions and protein duplications. Ideally the library is both large enough and minimally biased in order to produce a large fraction of possible variants. The product of this step can be considered a MISER library of plasmid pSAH060. **C)** A final cloning step moves the MISER library into a desired context – i.e. an expression plasmid, here pSAH063. Step C also allows for size-based exclusion of undesired protein variants by extraction from an agarose gel (Figure 1 and Figure S2).

**Figure S2.**

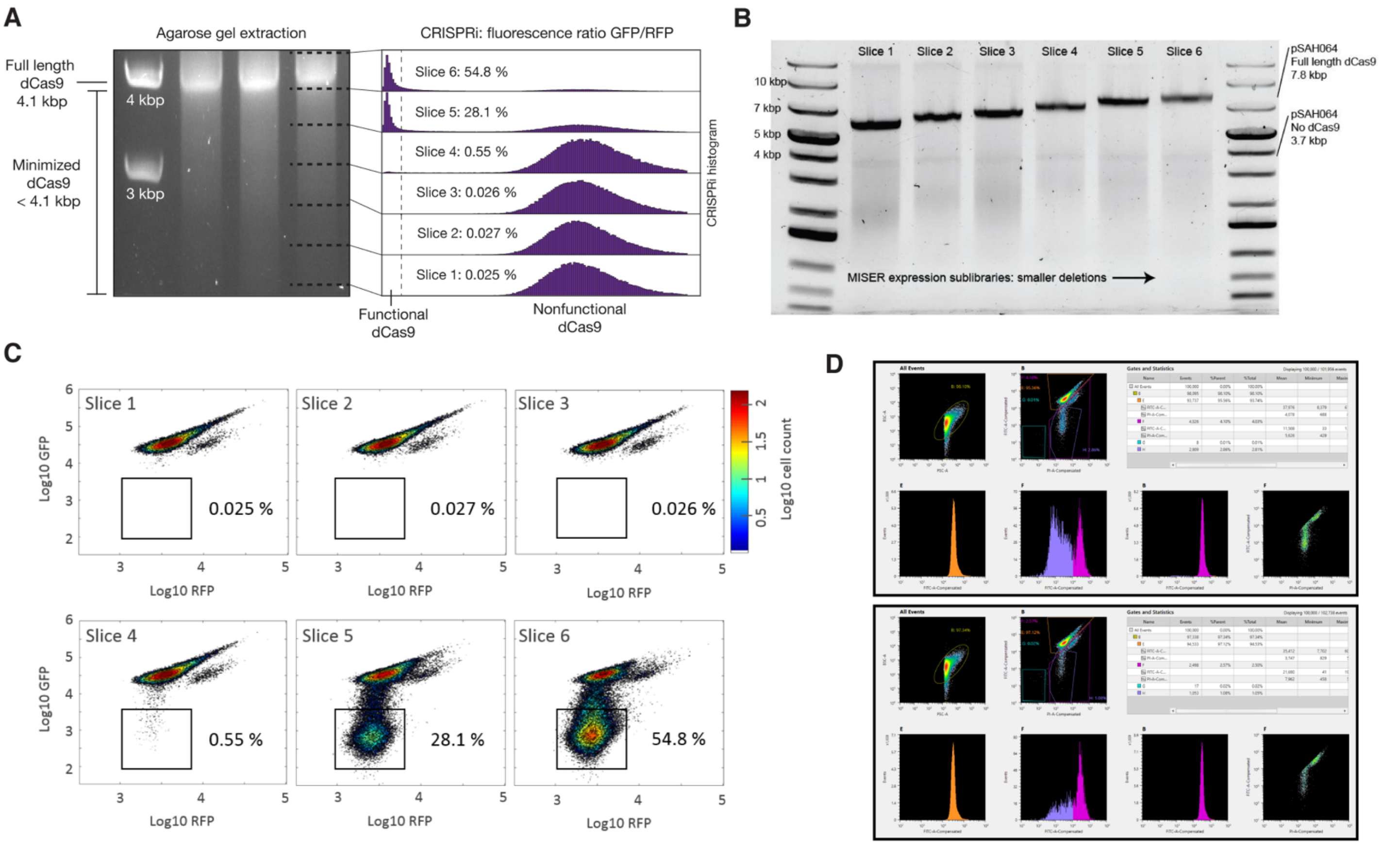

**<u>Figure S2:</u> Size exclusion and flow cytometry identify the range of dCas9 deletion sizes exhibiting *in vivo* transcriptional repression. A)** To empirically determine the size range of functional deletions, an agarose gel of the dCas9 MISER deletion library was sliced into six sub-libraries, independently cloned into expression vectors (B), and assayed for CRISPRi GFP repression via flow cytometry (C). Sublibrary Slice 4 was the most stringent library with detectable repression, with functional variants becoming more frequent in slices composed of smaller deletions as expected. **B)** The six gel slices in (A) were individually gel extracted and ligated into expression vector pSAH063, generating pSAH064 plasmids with dCas9 deletions. The resulting expression sub-libraries exhibit high precision in size ranges when assayed by agarose gel electrophoresis. **C)** Flow cytometry identifies Slice 4, 5, and 6 as expression sub-libraries containing functional dCas9 deletion variants. GFP repression CRISPRi was performed as described in Experimental Design. The region of phenotype defined as ‘functional’ is illustrated. The percent of functional hits is annotated. **D)** Screenshots from Sony Cell Sorter Software exemplifying the gating strategy, with upper panel showing full library sort and lower panel showing Slice 4. Gate H was used to sort cells containing repression-competent CE variants.

### Figure S3

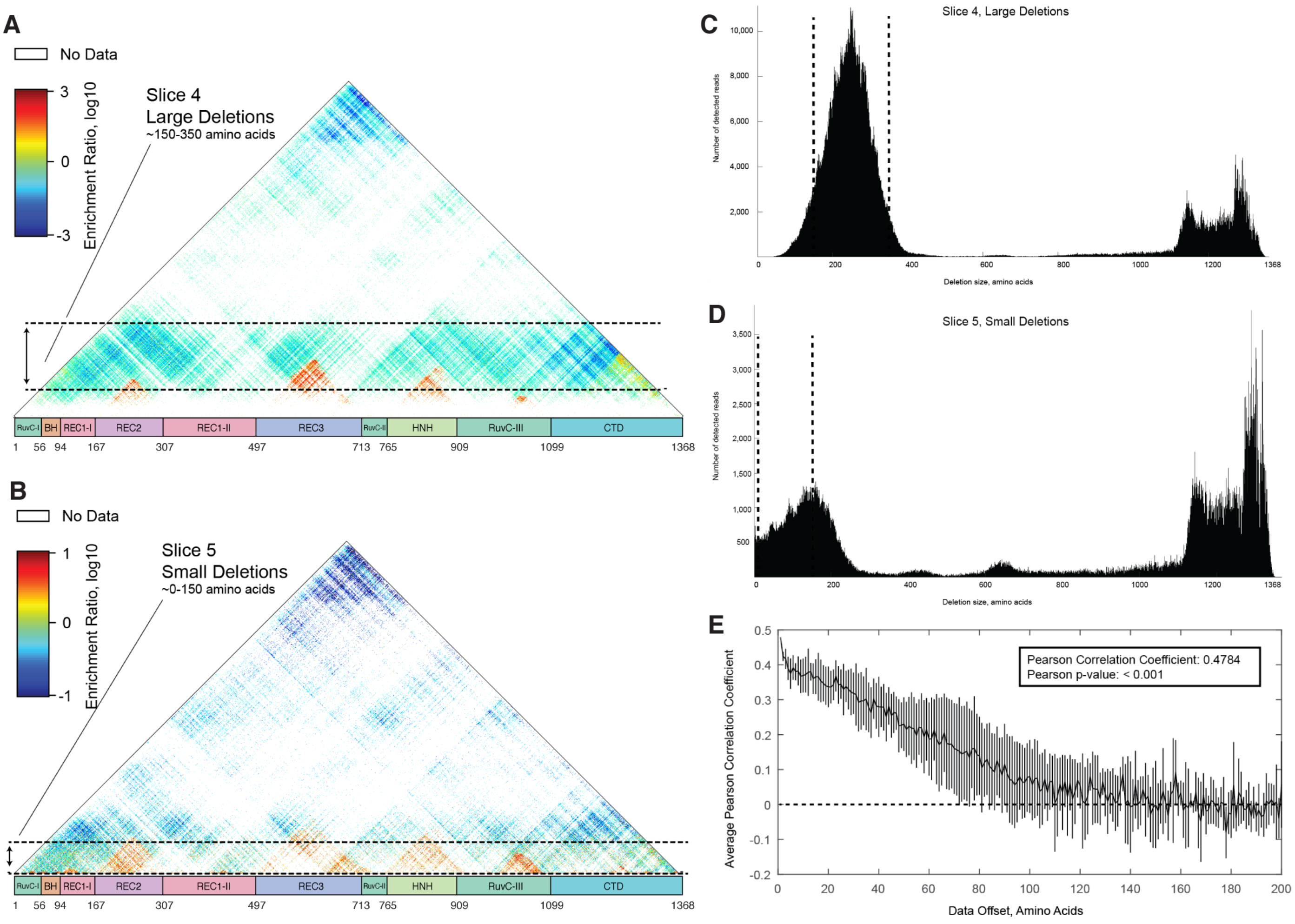

**<u>Figure S3:</u> Deep sequencing of the sublibraries of Slice 4 and Slice 5 reveal deletion regions throughout dCas9. A)** Raw enrichment map of Slice 4 sub-library. Each pixel represents a single deletion variant, whose start and end points are the axis intercepts when moving down and to the left or right, respectively, as described in the main text. Domain boundaries are labeled by amino acid number. The pixel color also denotes the degree of enrichment or loss following flow cytometry screening for transcriptional repression in vivo. Detailed calculations are described in the supplementary methods. Deletions corresponding to sizes within the gel slice are indicated by dashed lines. **B)** Raw enrichment map of Slice 5 sub-library, as in (A). Note the differing range of enrichment ratios. **C)** Histogram of deletion sizes in the naïve Slice 4 library. The hypothetical edges of the gel slice are indicated by dashed lines. **D)** Histogram of deletion sizes in the naïve Slice 5 library. The edges of the gel slice are indicated by dashed lines. **E)** Slices 4 and 5 independently replicate the same large functional deletion regions. The raw enrichment maps of Slice 4 and Slice 5 contain many of the same variants, and the Pearson correlation for these variants is highly significant ($p < 0.001$). Furthermore, this correlation is progressively lost if the two enrichment maps are shifted relative to one another. The line plots the mean of four additional Pearson correlations where the data array has been offset – either up, down, left, or right – by the indicated number of amino acids. This analysis verifies that the two enrichment maps independently identify large-scale regions of dCas9 which can be deleted and validates the apparent visual correspondence between maps A and B. Error bars, standard deviation.

**Figure S4.**

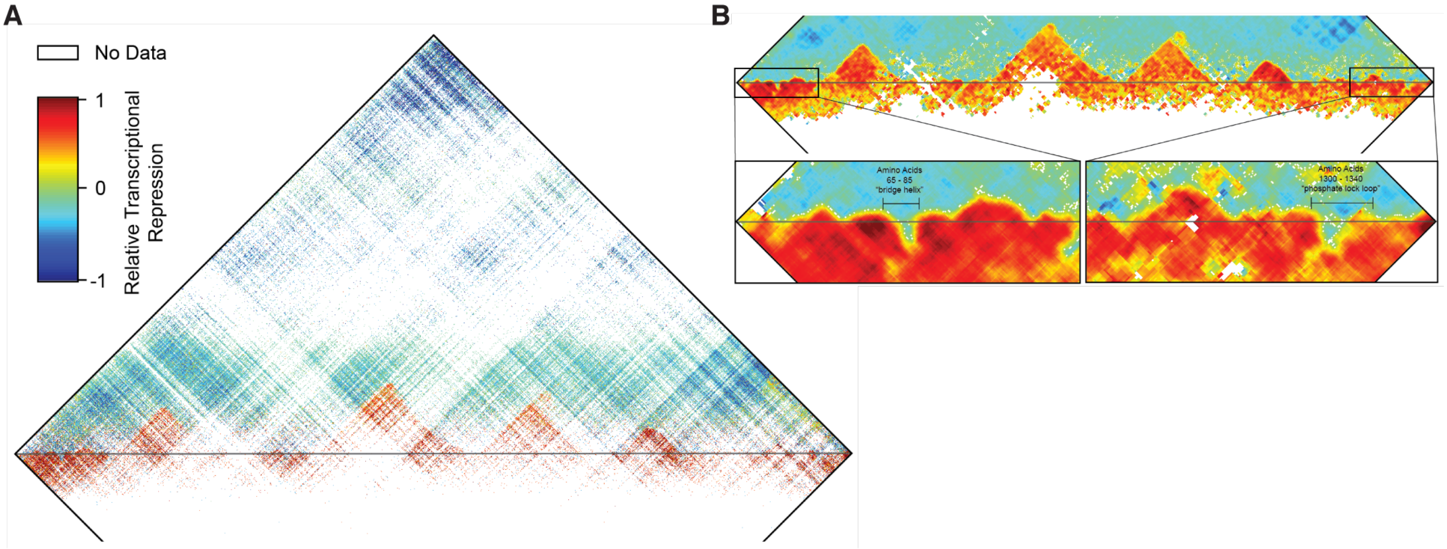

**<u>Figure S4:</u> Key elements of dCas9 secondary structure are revealed by the functional impact of small deletions and insertions. A)** The enrichment map of Figure 1C is presented in its entirety, including small duplications of dCas9 sequence. The horizontal grey line corresponds to the boundary between deletions (top) and tandem duplicate insertions (bottom). Note that in all cases a two amino acid MISER scar is also present (either Ala-Ser or Thr-Ser) which is not included in display or numbering. **B)** The combined enrichment map in (A) was interpolated to highlight the boundaries between functional and non- functional deletions, which are not clearly visible in the raw data. Pixels were replaced by the mean enrichment value of neighboring deletions/duplications, plus itself, in a square window 10 amino acids wide. Windows with fewer than five values were left white. Insets: The N- and C- terminal regions were particularly well resolved by this method, and elements of interest are annotated. The 'bridge helix' and 'phosphate lock loop' are two examples of secondary structure which strongly disallow small insertions.

**Figure S5.**

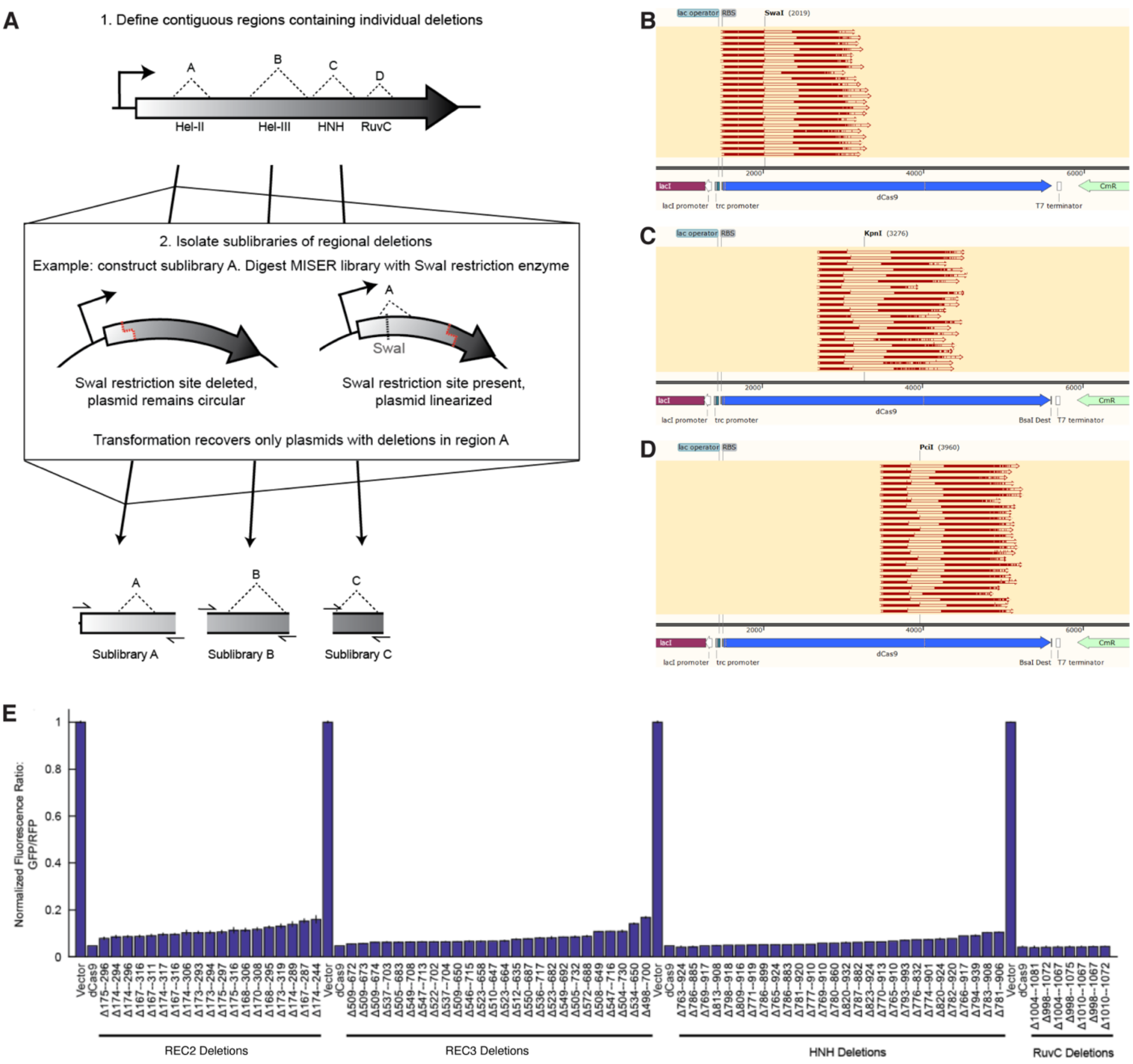

**<u>Figure S5:</u> MISER sublibraries composed of specific deletions can be generated by restriction digestion.** **A)** Digesting a MISER library with a restriction enzyme that has exactly one site within the plasmid will linearize the majority of plasmids, while plasmids with the site deleted will remain circular. This reaction can then be transformed in order to recover a sublibrary containing deletions from a specific region. **B)** For example, the restriction enzyme SwaI was used to isolate deletions in the REC2 region. The enzyme recognition site is shown mapped to the sequence of pSAH064, the dCas9 expression plasmid, illustrating the overlap with various sequenced deletions. **C)** The restriction enzyme KpnI was used to isolate deletions in the REC3 region, as in B. **D)** The restriction enzyme PciI was used to isolate deletions in the HNH region, as in B. **E**) Sublibraries containing regional individual deletion variants were re- transformed, and colonies were picked and assayed for CRISPRi activity. A subset of the most active clones was Sanger sequenced to identify the precise deletion. RuvC deletions could not be isolated by the sublibrary approach, and instead were cloned manually by PCR. Data are plotted as mean±SD from biological triplicates.

**Figure S6.**

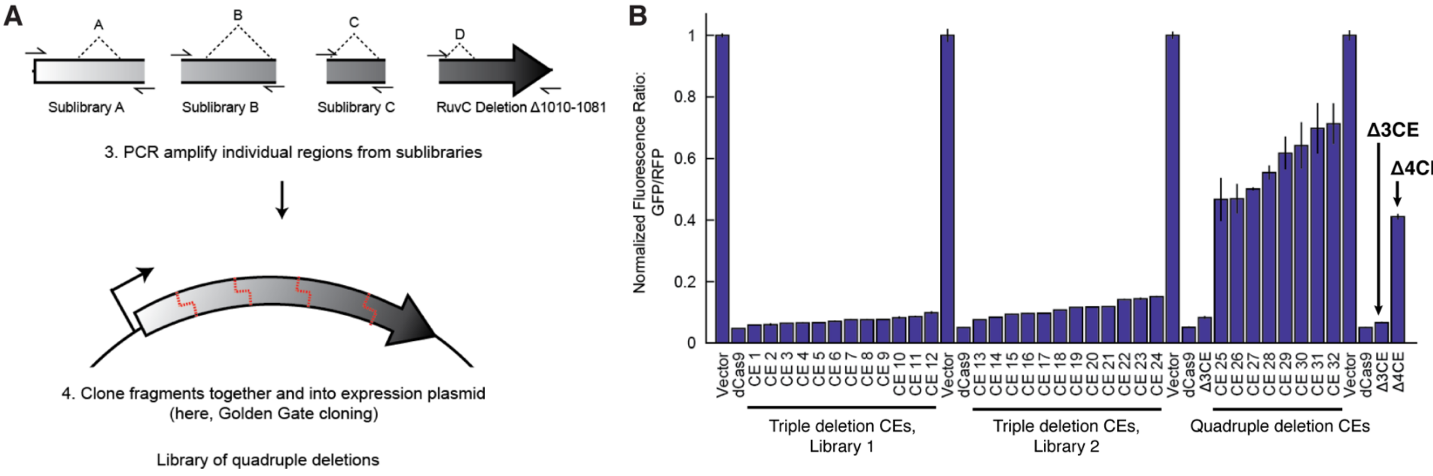

**<u>Figure S6:</u> Golden Gate Cloning builds libraries of CRISPR Effector (CE) variants with multiple deletions.** **A)** One highly functional RuvC deletion variant from Region D was PCR amplified, along with Sublibraries A, B, and C. PCR primers added Golden Gate compatible sticky ends, enabling Golden Gate cloning of individual fragments to form a library of CE deletion variants, Library 1. **B)** Flow cytometry was performed to isolate the most functional CE variants from the "stacked" library described in (A). All highly functional CE variants from Library 1 were found to lack REC2 deletions (sequences of CE variants selected for display on this plot can be found in Table S3). To verify this result, a second version of Sublibrary A was created, using a different strategy to isolate REC2 deletions as follows: the full MISER library was digested with the restriction enzyme BlpI, which cuts at amino acids 227-228 (instead of SwaI), and the resulting DNA was used directly as template for the PCR reaction (BlpI cuts pSAH064 three times and thus cannot be directly re-transformed to isolate the sublibrary). Library 2 thus contains all four deletion variants as in Library 1, except the sublibrary of REC2 deletions was entirely remade. However, once again functional CE variants isolated by FACS lacked REC2 deletions. The most functional variant in Library 2, CE 13, was named Δ3CE. Finally, to directly assay the effects of a REC2 deletion, the REC2 region of Δ3CE was replaced with a library of deletions from Sublibrary A. These quadruple deletion CE variants all exhibited vastly reduced CRISPRi activity compared to Δ3CE alone. The most functional variant assayed was named Δ4CE. Data are plotted as mean±SD from biological triplicates.

**Figure S7.**

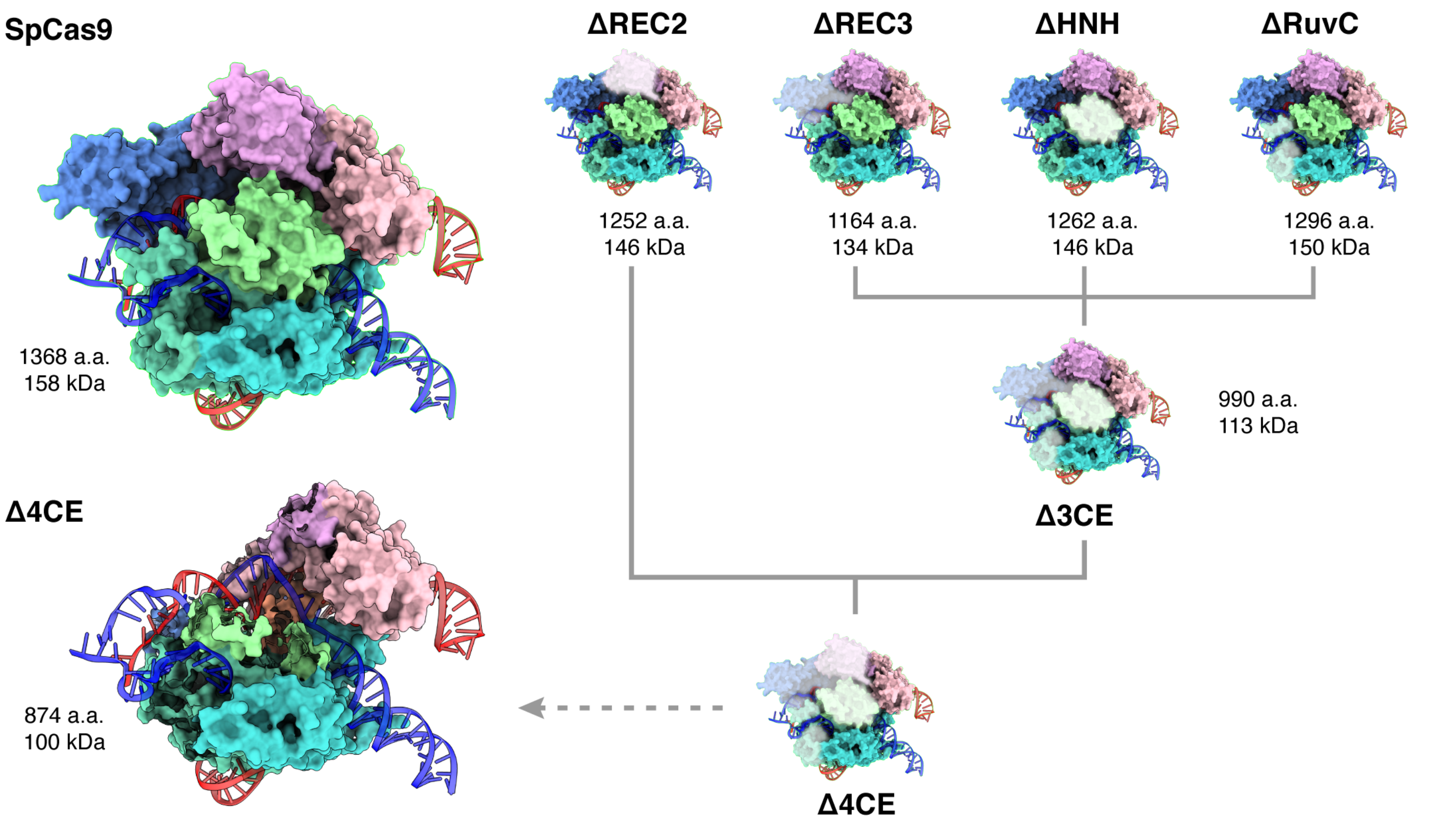

**Figure S7: 3D comparison of complete dCas9-sgRNA-dsDNA complex and modeled MISER constructs.** Model of SpCas9 complexed with sgRNA and dsDNA (PDB 5Y36), and MISER domain deletions overlaid. Δ3CE contains the REC3, HNH, and RuvC deletions, and Δ4CE contains the additional REC2 deletion, as described in Fig. 2 and S5. The Δ4CE model is shown with the domains corresponding to MISER deletions hidden. Molecular weights are calculated by the ExPASy ProtParam tool (https://web.expasy.org/protparam/).

**Figure S8.**

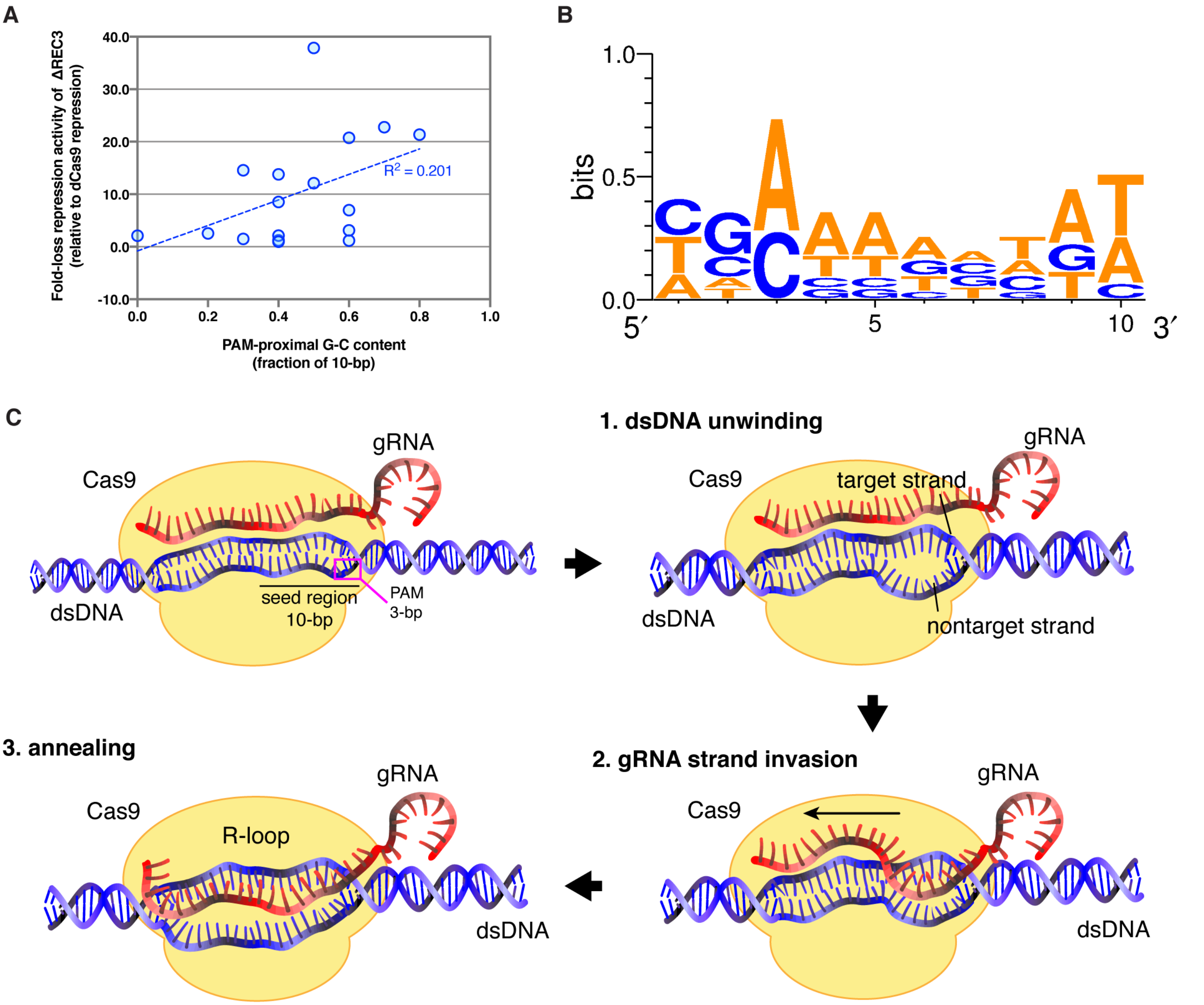

**<u>Figure S8:</u> Spacer sequence-dependent variability in repression activity of ΔREC3. A)** Plot showing fold-change in repression by ΔREC3 for different targets versus fraction of G-C content in seed region. Correlation between G-C content and repression is low and does not fully explain the variability in repression seen by the ΔREC3 construct across different target sequences. **B)** WebLogo showing spacer sequence variability for guides that exhibit at least a three-fold loss in repression by ΔREC3 compared to dCas9. **C)** Schematic showing the process of gRNA invasion into the dsDNA target leading to R-loop formation by Cas9. In Step 1, unwinding of the dsDNA double-helix is initiated at 1-2 bases adjacent to the PAM in the seed region, creating a destabilized region where the gRNA can invade, in Step 2. Hybridization of the gRNA to the target strand occurs in the seed region and proceeds in the PAM-distal direction (3'→5'), until the entire spacer sequence (~20bp) is annealed to the target strand, generating an RNA-DNA duplex called an R-loop (Step 3). RNA-DNA hybrid is shown as a 2-D representation for clarity instead of a helix.

**Figure S9.**

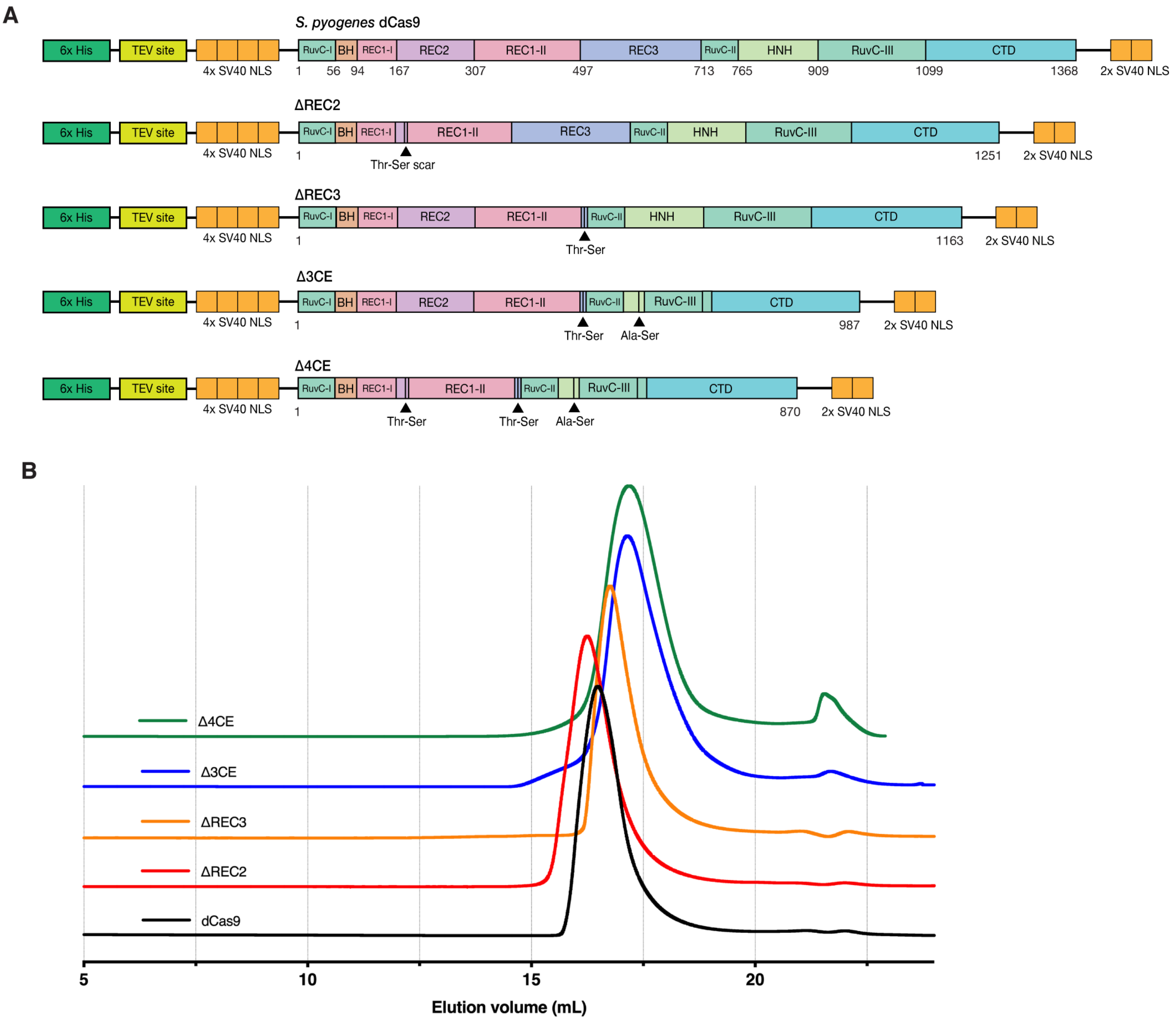

**<u>Figure S9</u>: Expression constructs and protein purification of MISER constructs. A)** Expression constructs for dCas9 containing MISER deletions and accompanying scars. All constructs were expressed using an IPTG-induced T7 promoter, and contain a N-terminal 6x His-tag, a TEV protease site, 4x SV40 NLS, and 2x SV40 NLS on the C-terminus. **B)** Size-exclusion chromatogram showing elution of all MISER constructs on a GE Superose 6 Increase column.

**Figure S10.**

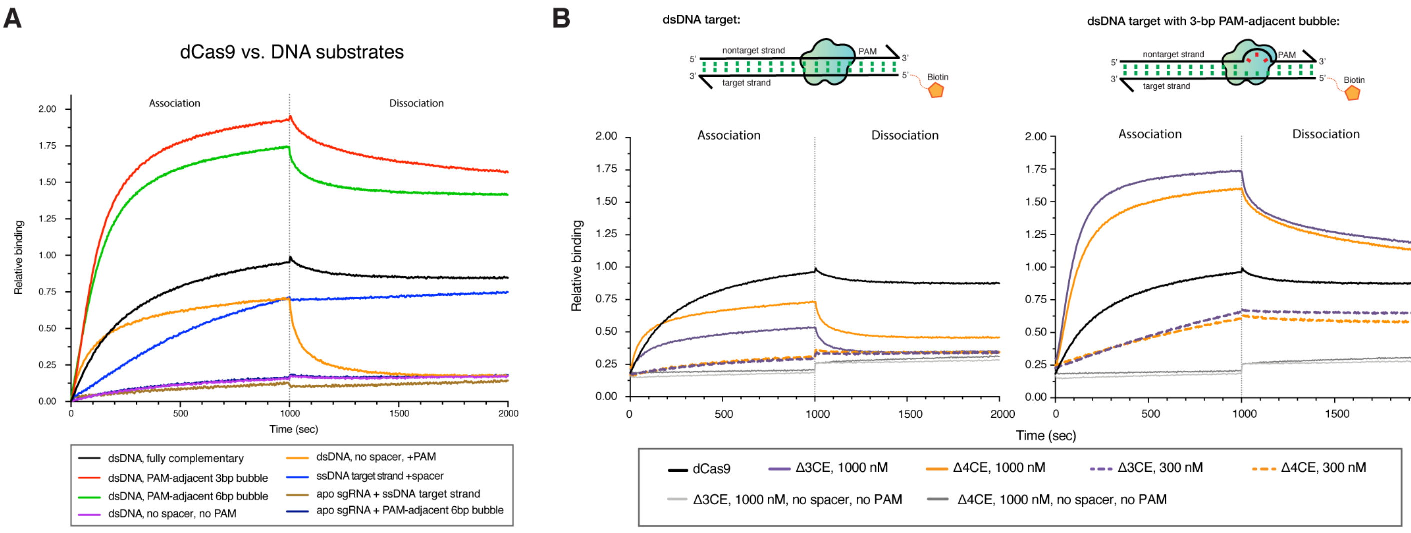

**<u>Figure S10:</u> Bio-layer interferometry (BLI) controls. A)** BLI experiments were performed by incubating immobilized dCas9 with dsDNA containing a target spacer but no PAM (orange trace). Transient PAM interactions have a significant contribution to the $k_{on}$ of association. The signal is lost immediately in the dissociation step, which suggests that the interaction is nonspecific. Conversely, incubation with a dsDNA containing no spacer and no PAM shows no signal (purple). **B)** BLI traces of Δ3CE and Δ4CE binding to dsDNA show that the relative binding is minimal at 300 nM, even with a 3-bp bubble in the seed region of the target (orange and purple). Subsequently a concentration of 1000 nM was used for these constructs. Light grey and dark grey traces represent Δ3CE and Δ4CE RNPs, respectively, against dsDNA without a spacer or PAM. All data shown are normalized to the maximum signal of dCas9 vs. fully complementary dsDNA target (black).

**Figure S11.**

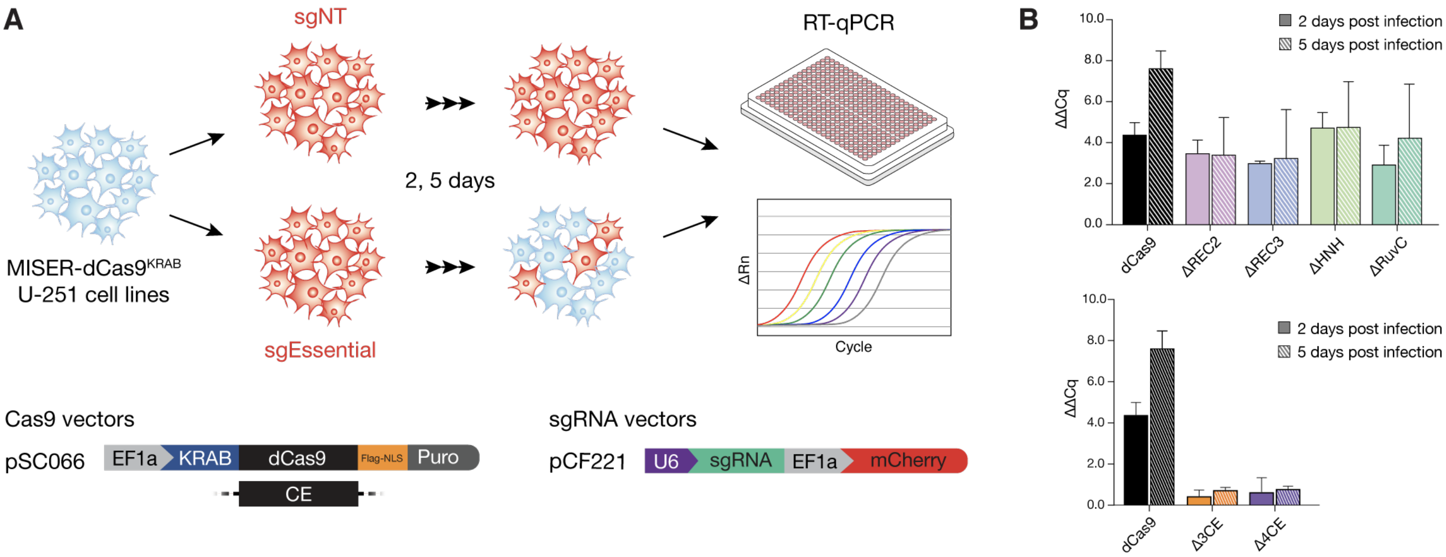

<u>**Figure S11:**</u> **Schematic of CRISPR interference (CRISPRi) based survival assay. A)** U-251 glioblastoma cells are stably transduced with lentiviral vectors (pSC066) expressing MISER-dCas9 or WT-dCas9 KRAB fusion proteins, followed by selection on puromycin. The various cell lines are then transduced with a secondary lentiviral vector (pCF221) expressing mCherry fluorescence protein and either sgRNAs targeting essential genes (sgPCNA) or non-targeting sgRNAs (sgNT) as controls. Cells are grown and harvested 2 and 5 days post-infection for RNA extraction, followed by RT-qPCR to quantitate transcription of targeted essential genes under MISER-KRAB repression. **B)** PCNA $\Delta\Delta C_q$ values from RT-qPCR at 2 (solid) and 5 (hatched) days post infection, calculated by subtracting target samples from sgNT samples. Values are plotted from biological duplicates as mean±S.D.

**Figure S12.**

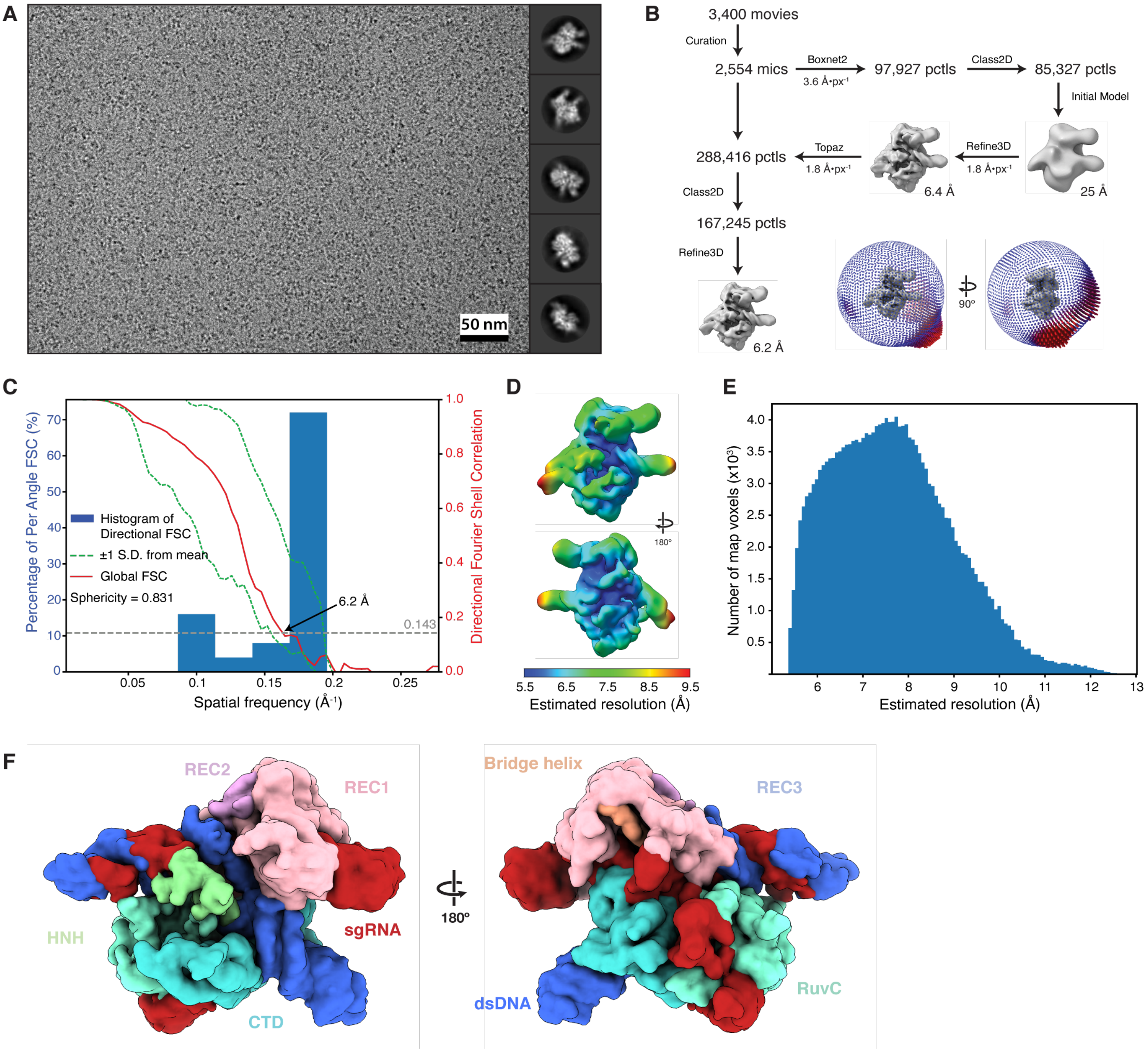

**<u>Figure S12:</u> Single-particle cryo-EM of the Δ4Cas9 ternary complex. A)** Exemplar micrograph at approximately 3 microns defocus with scale indicated and representative reference-free 2D class averages from the Topaz-picked particle set. Diameter of 2D mask is 150 Å in all averages. **B)** Single-particle reconstruction work-flow as described in methods and orientation distribution of the final reconstruction inset. (C) Directional FSC for final reconstruction. **D) and E)** Local resolution estimates calculated in RELION shown by coloration on the map and as a histogram, respectively. **F)** Density map of Δ4CE with putative domains segmented and colored according to their relative position within a 20 Å radius when overlaid on WT SpCas9 (PDB 5Y36).

**Table S1.**

| | **SpeI Insertion** | **NheI Insertion** |
|---|---|---|
| **Recombineering Oligo: Insertion Site 1** | AACACGTCCGTCCTAGAACTcgtctcatacgcaa<br>Accgcctctccccgcgcgttggcggtctcaatct<br>ATGactagtgataagaaatactcaataggcttag<br>ctatcggcacaaatagcgtcgggagacgGCAAGC<br>GGTACACTCAGATCAGTGTTGAGCGTAACCAAGT | AACACGTCCGTCCTAGAACTcgtctcatacgcaa<br>Accgcctctccccgcgcgttggcggtctcaatct<br>ATGgctagcgataagaaatactcaataggcttag<br>ctatcggcacaaatagcgtcgggagacgGCAAGC<br>GGTACACTCAGATCAGTGTTGAGCGTAACCAAGT |

**Table S1: Example Oligo Library Synthesis (OLS) oligonucleotides used in this study.** The full list of ordered oligonucleotides is available as ‘Auxiliary Supplementary Materials - Recombineering Oligonucleotides’. All oligonucleotides were ordered from Agilent Technologies, Inc. Oligos were designed to incorporate 45 and 47 bp of homology upstream or downstream of the insertion site, respectively (lowercase). Six bp were inserted between dCas9 codons, beginning after the target codon. The above example targets the start codon, ‘ATG’ (bold uppercase). These six bp consisted of recognition sequences for either the restriction enzyme SpeI or NheI (underlined). Flanking primer sequences allowed the amplification of the entire OLS library (italics) using oligonucleotides SAH_284 and SAH_285 (Table S6). Specific libraries of SpeI recombineering oligonucleotides or NheI recombineering oligonucleotides were amplified using forward primer SAH_284 and either SAH_286 or SAH_287 reverse primers, respectively. After amplification, these dsDNA products can be ‘matured’ by cleavage with the restriction enzyme BsmbI (bold lowercase), which cleaves internally of its recognition site, thus removing all non-homologous priming sequence from the recombineering template.

**Table S2.**

| Deletion | Δ3CE v1 | Δ3CE v2 | Δ3CE v3 | Δ3CE v4 | Δ3CE v5 | Δ3CE v6 | Δ3CE v17 | Δ3CE v21 | Δ3CE v22 | Δ3CE | Δ4CE |
|---|---|---|---|---|---|---|---|---|---|---|---|
| **REC2** | - | - | - | - | - | - | - | - | - | - | [180-297] |
| **REC3** | [511-716] | [498-699] | [500-688] | [497-700] | [501-664] | [512-721] | [509-650] | [508-649] | [508-646] | [503-708] | [503-708] |
| **HNH** | [813-909] | [813-908] | [811-898] | [786-882] | [804-893] | [809-916] | [776-923] | [768-900] | [786-923] | [792-897] | [792-897] |
| **RuvC** | [1010-1081] | [1010-1081] | [1010-1081] | [1010-1081] | [1010-1081] | [1010-1081] | [1010-1081] | [1010-1081] | [1010-1081] | [1010-1081] | [1010-1081] |

**Table S2: Deletions present in selected MISER variants. Indicated numbers represent the first and last amino acid deleted from the protein.**

**Table S3.**

| | Total Reads | Deletions Sequenced | Unique Deletions | Enriched Unique Deletions | De-enriched Unique Deletions |
|---|---|---|---|---|---|
| **Slice 4 Naïve** | 132,274,232 | 1,923,543 | 192,447 | | |
| **Slice 4 Sorted** | 140,589,968 | 1,960,138 | 25,948 | 19,618 | 6,330 |
| **Slice 5 Naïve** | 37,873,068 | 590,859 | 111,438 | | |
| **Slice 5 Sorted** | 35,016,326 | 290,947 | 51,462 | 31,794 | 19,668 |
| **Total** | 345,753,594 | 4,765,487 | 381,295 | 51,412 | 25,998 |

**Table S3: Statistics for deep sequencing of MISER libraries Slice 4 and Slice 5.**

**Table S4.**

| Gene | Distance from RBS (bp) | PAM-proximal 10bp sequence (5'-3') | PAM-proximal G-C fraction | Fold loss | Std. dev. |
|---|---|---|---|---|---|
| **GFP** | **38** | **AACAAGAATT-NGG** | **0.2** | **2.54** | **0.23** |
| RFP | 124 | TTAGCGGTCT-NGG | 0.5 | 37.84 | 3.78 |
| **GFP** | **130** | **ATAAATTTAA-NGG** | **0.0** | **2.11** | **0.01** |
| **GFP** | **174** | **TGACAAGTGT-NGG** | **0.4** | **1.23** | **0.02** |
| **GFP** | **196** | **TGAACACCAT-NGG** | **0.4** | **2.14** | **0.10** |
| **GFP** | **225** | **TCATGTGATC-NGG** | **0.4** | **0.96** | **0.05** |
| GFP | 262 | CCTTCGGGCA-NGG | 0.7 | 22.77 | 0.73 |
| **GFP** | **316** | **CGCGTCTTGT-NGG** | **0.6** | **1.18** | **0.06** |
| **GFP** | **355** | **CGATTAACAA-NGG** | **0.3** | **1.50** | **0.06** |
| RFP | 111 | TACCTTCGTA-NGG | 0.4 | 8.54 | 0.50 |
| RFP | 130 | TTCAGTTTAG-NGG | 0.3 | 14.56 | 0.77 |
| **RFP** | **165** | **CCCAAGCGAA-NGG** | **0.6** | **3.13** | **0.06** |
| RFP | 182 | CTGCGGGGAC-NGG | 0.8 | 21.35 | 0.71 |
| RFP | 197 | GGAACCGTAC-NGG | 0.6 | 6.98 | 0.23 |
| RFP | 208 | ACGTAAGCTT-NGG | 0.4 | 13.79 | 2.92 |
| RFP | 239 | CAGGTAGTCC-NGG | 0.6 | 20.74 | 4.25 |
| RFP | 248 | GGACAGTTTC-NGG | 0.5 | 12.10 | 0.60 |

**Table S4: gRNA target loci and G-C content dependence of ΔREC3 repression.** Spacer sequences highlighted in blue were used to generate the WebLogo in Figure S9A.

**Table S5.**

| **EMDB-22518** | |
|---|---|
| **Data Collection** | |
| Microscope | Talos Arctica |
| Magnification | 45,000 |
| Voltage (kV) | 200 |
| Detector | K3 |
| Electron exposure (e-/Å$^2$) | 60 |
| Defocus range (μm) | 1.5 to 3.8 |
| Pixel size (Å) | 0.45[a] |
| **Reconstruction** | |
| Symmetry imposed | C1 |
| Box size (pixels/Å) | 128/230 |
| Initial particle images (no.) | 288,416[b] |
| Final particle images (no.) | 167,245 |
| Map resolution (Å) | 6.2 |
| FSC threshold | 0.143 |
| Sharpening factor (Å$^2$) | -395 |
| Map resolution range (Å) | 5.5-9.5 |
| Sphericity | 0.831 |
| **Modeling** | |
| Method | Rigid-body |
| Initial Model | 5Y36 |
| CC | 0.73 |

[a]Super-resolution
[b]from picking with Topaz

**<u>Table S5:</u> Cryo-EM data collection & reconstruction statistics.**

**Table S6.**

| Oligo ID | Purpose | Sequence (5'-3') |
|---|---|---|
| SAH_284 | Recombineering amplification: universal forward | AACACGTCCGTCCTAGAACT |
| SAH_285 | Recombineering amplification: universal reverse | ACTTGGTTACGCTCAACACT |
| SAH_286 | Recombineering amplification: SpeI-specific reverse | GATCTGAGTGTACCGCTTGC |
| SAH_287 | Recombineering amplification: NheI-specific reverse | GATCGCCTAGACAACTCCTG |
| sgRNA-B9 | sgRNA for Cas9 RNP, used in BLI and cryo-EM | AGUCGGUGUCGACCCGGACCCAAAAUCUCGAUCUUUAUCGUUCAAUUUUAUUCCGAUCAGGCAAUAGUUGAACUUUUUCACCGUGGCUCAGCCACGAAAA |
| oAS081 | 5'-biotinylated ssDNA target for BLI, sgRNA-B9 | GCTCAATTTTGACAGCCCACCAGGCCCAGCTGTGGCTGATGGCATCCTTCCACTC |
| oAS003a | non-target ssDNA for BLI (complementary to oAS081) | GAGTGGAAGGATGCCATCAGCCACAGCTGGGCCTGGTGGGCTGTCAAAATTGAGC |
| oAS114 | 5'-biotinylated ssDNA non-target for BLI (no spacer, no PAM) | GTGTGCACACATGCAATAACATTGTGCACATGATACATTGCAATGACAATTAACC |
| oAS036 | non-target ssDNA for BLI (complementary to oAS081, 3-bp PAM-proximal bubble) | GAGTGGAAGGATGCCATCAGCCACAGCTGGGCCGATTGGGCTGTCAAAATTGAGC |
| oAS116 | unlabeled ssDNA target for BLI, sgRNA-B9. Used for cryo-EM RNP complex | GCTCAATTTTGACAGCCCACCAGGCCCAGCTGTGGCTGATGGCATCCTTCCACTC |
| sgNT-1 | Non-targeting gRNA for mammalian CRISPRi | GGCCAAACGTGCCCTGACGG |
| sgNT-2 | Non-targeting gRNA for mammalian CRISPRi | GCGATGGGGGGGTGGGTAGC |
| sgPCNA-i1 | *PCNA* targeting gRNA for mammalian CRISPRi | GGGGCGAACGTCGCGACGAC |
| sgPCNA-i2 | *PCNA* targeting gRNA for mammalian CRISPRi | GGCGTGGTGACGTCGCAACG |
| sgPCNA-i3 | *PCNA* targeting gRNA for mammalian CRISPRi | GCGCTCCCGCCAAGCACCGG |
| sgPCNA-i4 | *PCNA* targeting gRNA for mammalian CRISPRi | GAAGCGCTCCCGCCAAGCAC |
| sgPCNA-i5 | *PCNA* targeting gRNA for mammalian CRISPRi | GCCCGGCCCGCCTGCACCTC |
| sgPCNA-i6 | *PCNA* targeting gRNA for mammalian CRISPRi | GCGGACGCGGCGGCATTAAA |
| sgPCNA-i10 | *PCNA* targeting gRNA for mammalian CRISPRi | GGCCATCCGCGCCTTCTCAT |
| sgRPA1-i1 | *RPA* targeting gRNA for mammalian CRISPRi | GGGAAGCTGGAGCTGTTGCG |
| sgRPA1-i2 | *RPA* targeting gRNA for mammalian CRISPRi | GGCGACGGGGGATGAACGCG |
| sgRPA1-i3 | *RPA* targeting gRNA for mammalian CRISPRi | GTGCGCAGCGCGCGGGACCC |
| sgRPA1-i4 | *RPA* targeting gRNA for mammalian CRISPRi | GTGAGCCGCGCGCACGTCGG |
| sgRPA1-i5 | *RPA* targeting gRNA for mammalian CRISPRi | GGCGGTGCGCGCAACTTCTC |
| sgRPA1-i8 | *RPA* targeting gRNA for mammalian CRISPRi | GCGAGCCTCGCGGAGTAGAG |
| sgRPA1-i9 | *RPA* targeting gRNA for mammalian CRISPRi | GCCGCGCGCTGCGCAGTTAT |
| oAS085 | Forward primer for *RPA1* cDNA reverse transcription, set 1 | GCAGTTGGAGTGAAGATTGG |
| oAS086 | Reverse primer for RPA1 cDNA RT, set 1 | CACTTGGACTGGTAAGGAGT |
| oAS087 | Forward primer for *RPA1* cDNA RT, set 2 | CCGAGCTACAGCTTTCAATG |
| oAS088 | Reverse primer for RPA1 cDNA RT, set 2 | GCAGATCCCGATGATGTCTA |
| oAS089 | Forward primer for *PCNA* cDNA RT, set 1 | ACTCAAGGACCTCATCAACG |
| oAS091 | Reverse primer for PCNA cDNA RT, set 1 | TGAACCTCACCAGTATGTCC |
| oAS090 | Forward primer for *PCNA* cDNA RT, set 2 | CGTTATCTTCGGCCCTTAGT |
| oAS092 | Reverse primer for PCNA cDNA RT, set 2 | CGTGCAAATTCACCAGAAGG |
| oAS117 | Forward primer for *GAPDH* RT | TCAAGGCTGAGAACGGGAAG |
| oAS118 | Reverse primer for *GAPDH* cDNA RT | TGGACTCCACGACGTACTCA |
| oAS034 | Forward primer for cloning dCas9 and MISER constructs into expression vector | GGTATCAACTTTTCGTTTCTT |
| oAS035 | Reverse primer for cloning dCas9 and MISER constructs into expression vector | CAAAGCCCGAAAGGAAG |

**<u>Table S6:</u> Oligonucleotides used in this study.**
